# The ARF regulatory GTPase in *Giardia intestinalis* is associated with vesicle formation and membrane fusion machinery at non-canonical endosomal compartments

**DOI:** 10.1101/2025.10.20.683495

**Authors:** Shweta V. Pipaliya, Emily M. Kingdon, Corina D. Wirdnam, Erina A. Balmer, Richard A. Kahn, Joel B. Dacks, Carmen Faso

**Author notes:** Co-corresponding author (CF), (JBD). Equally contributing authors (Shweta V. Pipaliya, Emily M. Kingdon).

## Abstract

The intestinal parasite *Giardia intestinalis*, the causative agent of the globally distributed diarrheal disease Giardiasis, is one of the few genetically tractable members of the phylum Fornicata, which includes both parasitic and free-living species. The diversity of membrane traffic machinery in this lineage is of special interest in relation to the evolution of parasitism and the emergence of specialized organelles as possible adaptations to parasitism.

Here, we performed a functional characterization of the ARF family of regulatory GTPases and their regulators in Giardia traffic, including three ARF paralogues, one ARF GAP and one ARF GEF. Using a combination of bioinformatic tools, protein network discovery and validation, and confocal light microscopy, we show that the Giardia ARF complement studied here is robustly associated with peripheral endocytic compartments (PECs), essential feeding organelles that are unique to Giardia parasites. Intersection of the interactomes of this ARF complement with previously published data for PECs-proteins (including membrane adaptors, predicted retromer subunits and SNAREs), revealed a complex crosstalk between the ARF complement and several membrane traffic processes.

## Introduction

The ARF family is a collection of small, regulatory GTPases that are involved in a wide range of cellular functions. This family includes both the ARF proteins, which typically share >65% primary sequence identity, and the ARF-like (ARL) proteins that are more divergent, sharing typically ∼40-60% identity. Herein we focus exclusively on the ARFs, which are crucial regulators of membrane traffic and organelle architecture in eukaryotic cells, operating through a tightly controlled cycle of ARF GEF (guanine nucleotide exchange factor)-mediated GTP binding and ARF GAP (GTPase activating protein)-mediated GTP hydrolysis [1]. ARF activation is tightly linked to their reversible membrane association because of changes in the conformation of the myristoylated amphipathic helix at their N-termini. An ARF signaling “module” is often portrayed as the sequential actions of a specific ARF GEF acting on a specific ARF GTPase, which upon activation recruits specific protein effectors to the membrane surface and subsequent termination by a specific ARF GAP. However, it is increasingly clear that such systems are more complex as both the GEFs and GAPs can be regulated by other interactions and the specificities of each may be influenced as a result. In mammals ARF proteins regulate the recruitment of cytosolic coat complexes to membranes at several sites in both anterograde and retrograde membrane traffic, though are most often linked to actions at the Golgi complex and trans-Golgi network (TGN). Subsequent research has revealed that ARF proteins have a broader range of roles, including involvement in phospholipid metabolism and signaling, cytoskeletal organization, lipid droplet management, and non-vesicular lipid transport [1–8].

In humans, there are 5 Arf proteins (Arfs 1,3,4,5, and 6), with Arfs 1-5 are derived from an ancestral Arf1 progenitor and Arf2 lost in primates. The ancestor that gave rise to all eukaryotes (i.e. the Last Eukaryotic Common Ancestor or LECA) is inferred to have possessed orthologues of Arf1 and Arf6. Similarly, there are 16 known ARF GEFs [9] and 28 ARF GAPs [10] in humans, having been expanded from an complement of 3 ARF GEFs (BIG, GBF, and Cytohesin) and at least seven ARF GAPs inferred to have been present in the LECA [9,11,12]. Knowing this ancestral complement is key to understanding of the ancient evolution of the eukaryotic membrane traffic system. It also provides a baseline from which to deduce the evolution across different eukaryotic lineages, whether expansion as in humans or reduction in organisms with reduced subcellular organization and/or specialized traffic routes.

The phylum Fornicata is one such lineage. It consists of unicellular species, both free-living and parasitic [13], including the genetically tractable species *Giardia intestinalis* (syn. *lamblia*, *duodenalis*), an anthropozoonotic parasite of the mammalian small intestine, of global medical importance. Giardia is also a compelling eukaryotic model because of its reduced protein traffic-related organellar and molecular complements [14].

*G. intestinalis* alternates between a flagellated (trophozoite) and cyst forms. The former colonizes the small intestine and feeds on luminal content through peripheral endocytic compartments (PECs) [15–17] In contrast, the cystic form allows for parasite survival outside a host and, following ingestion, initiates a new infection cycle. Trophozoites differentiate to cysts during a process called encystation, during which dedicated organelles called encystation-specific vesicles (ESVs) mediate traffic of cyst wall material [18–20].

While Giardia lacks many of the typical characteristics of membrane traffic [21], it presents with a basal endomembrane system consisting of only five membrane-bound compartments – one of which is an extensive endoplasmic reticulum (ER). Most distinctively, Giardia is missing the conventional Golgi apparatus and related cisternae, as well as genuine endosomes and mitochondria [21–23]. The membrane traffic system of Giardia has distinctly evolved, stemming from the adaptation to the intestinal lumen environment of the host [15,16]. At the proliferating trophozoite stage, Giardia have an endomembrane system, consisting of four organelles: (1) the nuclear envelope, (2) the ER, (3) mitosomes, and (4) the peripheral endocytic compartments (PECs).

Giardia employ PECs, previously referred to as peripheral vacuoles (PVs), as the only entry port for fluids. These organelles acidify and presumably serve as digestive compartments with the capability for sorting after processing [24] that is suggestive of a functional similarity between PECs, endosomes and lysosomes. The method of initiation for contact between the PEC and cell membrane is not yet understood. There is a growing body of evidence which points to PECs having a role in the release of potentially virulent proteins through unconventional protein secretion (UPS) routes [25]. There have been four UPS pathways determined to date, and it has been theorized that there is a linkage between PECs and UPS pathways [25,26].

The Fornicata as a whole possesses the characteristic of the ARF regulatory system reduced from the LECA complement early in the Fornicate lineage and prior to any shift to parasitism [12]. Comparative genomic and phylogenetic analyses have shown that Giardia possesses three ARF1 genes, one apparently more canonical in sequence and two more diverged ARF1 paralogues, ARF1FA and ARF1FB, which are unique to the fornicate lineage. No Arf6 has been reported in members of the Fornicata. The ARF regulatory system is simplified compared to that of the LECA, with a reduced complement of ARF GEFs (with only BIG and Cytohesin orthologs) and ARF GAPs (only SMAP, ARFGAP1, and AGFG orthologs). Because this complement appears to have been reached by the time of diversification in the Fornicata, it pre-dates the transition to parasitism in this lineage.

While the protein complement in Giardia is known, the cell biological roles of the Arf paralogues are poorly understood at best. The canonical ARF1 in Giardia localizes to PECs and ESVs in Giardia trophozoites and cysts, respectively and proposed to be necessary for transport and loading of cyst wall material during encystation [11]. Interference with the ability of ARF1 to function in Giardia leads to defective cyst wall formation [26]. In contrast, the ARF1 paralogs are hypothesized to assist in clathrin-mediated bulk flow processes for cargo internalization, potentially associating with the PECs and tubular ER [27].

This paucity of information is particularly glaring given the well-known paradigm of Arf1 activity at the Golgi body and the highly divergent nature of the Giardia endomembrane system, including the apparent lack of a Golgi body and the enigmatic nature of the endosomally derived PECs. To address this knowledge gap, we embarked on a systematic investigation of the evolution, the subcellular localization and the protein-protein interactions of *Gi*ARF1, *Gi*ARF1FA, *Gi*ARF1FB1 in Giardia trophozoites and their regulatory partners *Gi*ARFGAP1 and *Gi*CYTHa.

## Results

### Expanded molecular phylogeny of ARFs, GAPs and GEFs in BaSks and Anaeramoeba

Since the last determination of the Arf regulatory system complement across the Fornicata, several new and deeply divergent related organisms have been discovered and their genomes sequenced. This includes members of the BaSk clade (*Barthelona* sp. PAP020, *Barthelona* sp. PCE, *Skoliomonas litria*, *Skoliomonas* sp. GEMRC, *Skoliomonas* sp. RCL) which are basal to the Fornicata and Anaeramoeba (*Anaeramoeba flamelloides, Anaeramoeba ignava*) [27] which are sister to the parabasalids [28], another major group of the Metamonada supergroup to which the Fornicata belongs [29]. In order to better detail the reduction of the Arf regulatory system from the LECA complement to that seen in Fornicata, undertook a molecular evolutionary investigation in these new organismal genomes.

ARF1 similarity searching was performed with an HMM of eukaryotic sequences and identified copies of ARF1 in all sampled genomes. Surprisingly, ARF6 searches revealed candidate ARF6 orthologues in all new genomes except for *Skoliomonas* sp. RCL and *Skoliomonas litria*. Phylogenetic analyses of all candidate Arf1 and Arf6 proteins together were unresolved (Supplementary Figures 1-3). Therefore we analyzed the candidates separately against a previously resolved backbone dataset [30]. When ARF1 candidates were analyzed, they were resolved with high support as excluded from the Arf6 clade and within the Arf 1 clade without strong backbone support separating them from Arf1 sequences from other eukaryotes (Figure 1A). Similarly, the candidate ARF6 sequences were resolved with high support as excluded from the ARF1 clade and within the clade of ARF6 (Figure 1B).

**Figure 1.**
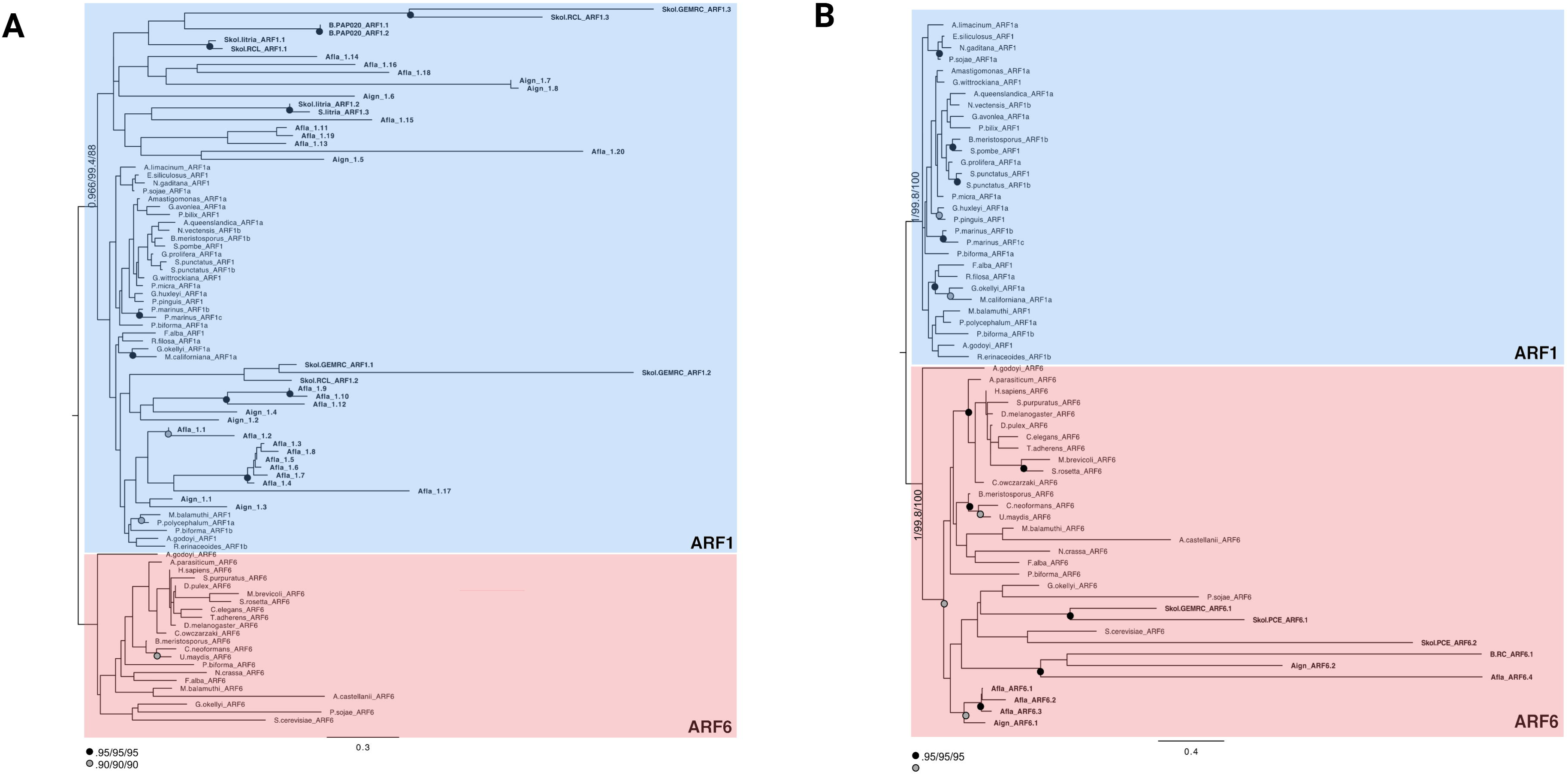
Phylogenetic investigation assessing the classification of ARF homologues in the BaSks and *Anaeramoeba*. (**A**) Classified ARF1 homologues in BaSks and *Anaeramoba* against an ARF backbone [30]. Phylogenetic analysis robustly supports the exclusion of the ARF1 sequences from the ARF6 clade. In this, values for supported nodes (MrBayes/SH-aLRT/IQ-TREE-2) have been replaced by symbols: black circles: >.95/95/95, grey circles: >.90/90/90. Node support values are overlaid onto IQ-TREE-2 tree topology. (**B**) Classified ARF6 homologues in BaSks and *Anaeramoba* against an ARF backbone [30]. Phylogenetic analysis robustly supports the exclusion of the ARF6 sequences from the ARF61clade. In this, values for supported nodes (MrBayes/SH-aLRT/IQ-TREE-2) have been replaced by symbols: black circles: >.95/95/95, grey circles: >.90/90/90. Node support values are overlaid onto IQ-TREE-2 tree topology.

Of the seven known ancestral ARF GAPs (SMAP, ArfGAP1, ArfGAP 2/3, ACAP, AGFG, ArfGAP_C2, and ADAP), we previously identified consistent conservation of only three in members of the Fornicata [11,12]; SMAP, AGFG and ARFGAP1. ARFGAP1 and 2 were both present in LECA but are often challenging to resolve by homology searching [11]. ArfGAP searches were performed using an HMM of the ArfGAP domain in eukaryotes. In all genomes, except *Skoliomonas* sp. GEMRC, SMAP, and an unresolvable ArfGAP1 *vs.* 2 protein were identified. In all BaSk genomes, AGFG was also detected.

From the LECA complement of GBF1, BIG, and cytohesin ARF GEFs appears to have been lost in the Fornicata [11]. Our searches into the BaSk and Anaeramoeba genomes were performed using an HMM of the Sec7/ArfGEF domain from diverse eukaryotes. Initial characterization of the hits shows the presence of cytohesin in *Barthelonids*, and *Anaeramoeba* and BIG in all but *Barthelona* sp. PAP020. GBF1 was identified only in the *Anaeramoeba* species.

The results obtained from the *Anaeramoeba* genomes are consistent with the close relative, pre-fornicate conclusions previously reported [9]. We saw positive hits for ARF1 and ARF6, which predates these losses. The presence of a canonical ARF1 within all these new genomes is unsurprising as ARF1 has, so far, not been lost in any investigated eukaryotic lineage [31]. Additionally, we saw cytohesin, BIG, and GBF1 within this group as well which predates the reduction of the ARF regulatory system in fornicates. We did, however, see a loss of the LECA AGFG and ArfGAP_C2 within the *Anaeramoeba* species. Within the new BaSk clade, we identified consistency in the presence of ARF1, as well as ARF6 in all species except for *Skoliomonas* sp. RCL. This does, however, make sense, as within the Skoliomonas group, *Skoliomonas* sp. RCL diverged prior to the other two species, so this is likely an independent loss. However, with the limited number of BaSk genomes, it is difficult to conclusively determine if this loss is limited to *Skoliomonas* sp. RCL. We do observe a consistent maintenance of AGFG, SMAP, and ArfGAP 2/3 within the BaSk clade, with a loss of and ArfGAP 2/3 in only the *Skoliomonas* sp. GEMRC genome. The consistent presence of ARF1 across all newly released and previously studied genomes reinforces its status as a consistently retained eukaryotic protein, whereas the variable presence of ARF6, ArfGEFs, and ArfGAPs displays patterns of lineage-specific retention and loss.

### In silico domain survey of GiARF1, GiARF1FA and GiARF1FB1

Having better defined the points of reduction and retention of the Arf regulatory system in the lineages leading to Giardia, we turned to focus our analysis on the Giardia Arf paralogues specifically. The three *Giardia* ARF paralogues previously identified [12] were subject to *in silico* sequence comparisons to assess residue conservation responsible for ARF GTPase enzymatic and effector binding activities. Overall, *Gi*ARF1 and *Gi*ARF1FA bear a high degree of sequence similarity to one another (Figure 2), while *Gi*ARF1FB1 is overall more divergent in its amino acid composition in comparison to the other two paralogues (Figure 2). Nonetheless, all three retain motifs that are characteristic of ARFs. These include an amphipathic alpha helix at their N-termini, and a glycine at position two that is predicted to be N-myristoylated [32,33] (Figure 2). Both the alpha helix and myristoylated N-terminal glycine were shown to be essential for regulated membrane association in mammalian cells [34].

**Figure 2.**
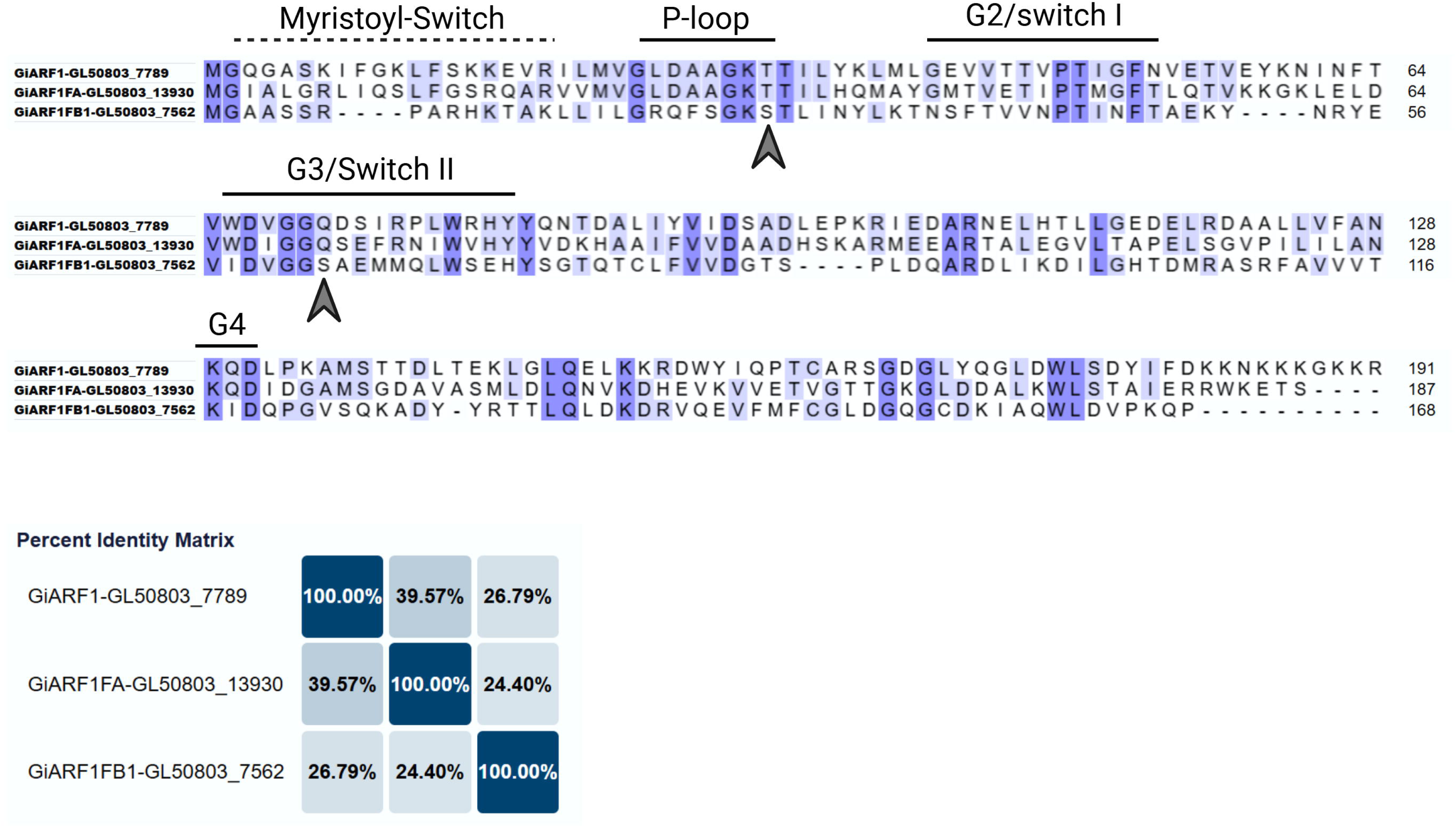
Sequence alignment of the three *Giardia intestinalis* AWB ARF1 paralogues for biochemical activity assessment. All three *Giardia* ARF1 sequences were examined to determine whether canonical motifs for myristoylation, Mg^2+^, GTP, GDP, and regulatory protein binding were present. All three ARFs contained the N-terminal myristate switch with a conserved glycine residue in position two necessary for membrane recruitment. The three paralogues were also characterized by the presence of four G-boxes (G1/P-loop, G2, G3, and G4) which coordinate GTP and GDP-binding. The P-loop and G3 also contain residues for Mg^2+^ and GAP binding, T/S31 and Q71, respectively (arrowheads). Overall, all three ARFs are predicted to retain canonical GTPase activity, but *Gi*ARF1FB1 is most diverged in its sequence composition compared to *Gi*ARF1 and *Gi*ARF1FA, which share significant similarities. Motif predictions were performed according to [36]. Alignment was generated using UniProt ClustalO.

Small, regulatory GTPases are characterized by the presence of highly conserved signature motifs that are used for guanine-nucleotide binding. These are termed G-boxes, and generally, five of them are present (G1-G5) [35,36]. The *Giardia* ARF paralogues retain four out of five of the G-boxes directly involved in GXP binding and therefore are predicted to do so. These are G1, also known as P-loop (GxxxxGK(T/S)T), G2 (PT), G3 (WDVGGQ), and G4 (NKxD) (Figure 2). The P-loop contains either a threonine or serine residue in position 31, necessary for Mg^2+^ binding to increase GTP-binding affinity [37,38] (indicated with the first blue arrow in Figure 2). G3 contains glutamine in position 71 and is the catalytic site for GTP hydrolysis [39] (indicated with the second blue arrow in Figure 2). Both *Gi*ARF1 and *Gi*ARF1FA contain a glutamine at this position, whereas *Gi*ARF1FB1 encodes a serine. This change is present in other ARF family members (e.g., ARL13) and is linked to differences in nucleotide binding and hydrolysis properties (Figure 2). Regulatory GTPases also contain sites for effector binding, termed switch I and switch II (that overlap with the G2 and G4 motifs), which are altered in conformation upon exchange of GDP for GTP resulting in increased affinity for binding partners [36].

### ARF1 localizes to the cortical PECs in vegetative trophozoites

To begin to assess the predicted cellular functions of the ARFs and regulators in Giardia trophozoites their cellular localizations were determined. Transgenic parasites expressing *Gi*ARF1 tagged with a C-terminal HA epitope (*Gi*ARF1-HA) were subjected to immunofluorescence microscopy (Figure 3). At least 150 cells were analysed, and 88% of the population expressed the *Gi*ARF1-HA variant (Supplementary Figure 4A-B). *Gi*ARF1-HA was found to accumulate distinctly at the cell periphery. Double labeling witha non-selective, soluble marker for PECs (Dextran-TxR) [40] revealed extensive overlap between the GiARF1-HA and PECs in the cortical region of the cell (Figure 3A). Co-localization analysis of *Gi*ARF1-HA and Dextran-TxR signal shows considerable signal overlap, however, only in regions bearing cortical PECs but not the bare zone (Figure 3C and D). Closer investigation of regions of interest (ROI) with the two PEC populations confirmed minimal, even complete absence, of *Gi*ARF1-HA at the bare zone (Figure 3C). Quantitative assessment of protein localization was performed through signal overlap analyses with at least ten cells to determine whether observed trends were uniformly present across multiple sampling points. Pearson’s coefficient values (R) in whole-image analyses ranged between 0.56 to 0.83 (average of 0.75), which suggested a high positive correlation between the two channels (Figure 3A, B and E; Supplementary Table 6). This was further corroborated by high M1 and M2 coefficient values, another metric of co-localization that measures the degree of pixel overlap from channel A to channel B and *vice versa* [41]. Average M1 and M2 values were 0.99 and 0.987, respectively (Figure 3B and E; Supplementary Table 6). A high degree of confidence in all three coefficients was confirmed by an average Costes’ p-value of 1, suggesting that the computed R, M1, and M2 coefficient values are not randomized. A similar positive correlative trend was observed with ROI2 analyses of the cortical PECs with an average R of 0.46, M1 and M2 of 0.99 and 1.00, respectively, and Costes’ p-value of 1 (Figure 3D and E; Supplementary Table 6). Statistical analyses confirmed a lack of co-localization between *Gi*ARF1-HA and Dextran-TxR in the bare zone region (ROI1), as negative or extremely low R values were observed across all sampled trophozoites. Although M1 and M2 coefficients are high, the Costes’ p-value is well below the required 0.95 threshold of confidence, suggesting that signal overlap between these two channels is unlikely (Figure 3C and E; Supplementary Table 6).

**Figure 3.**
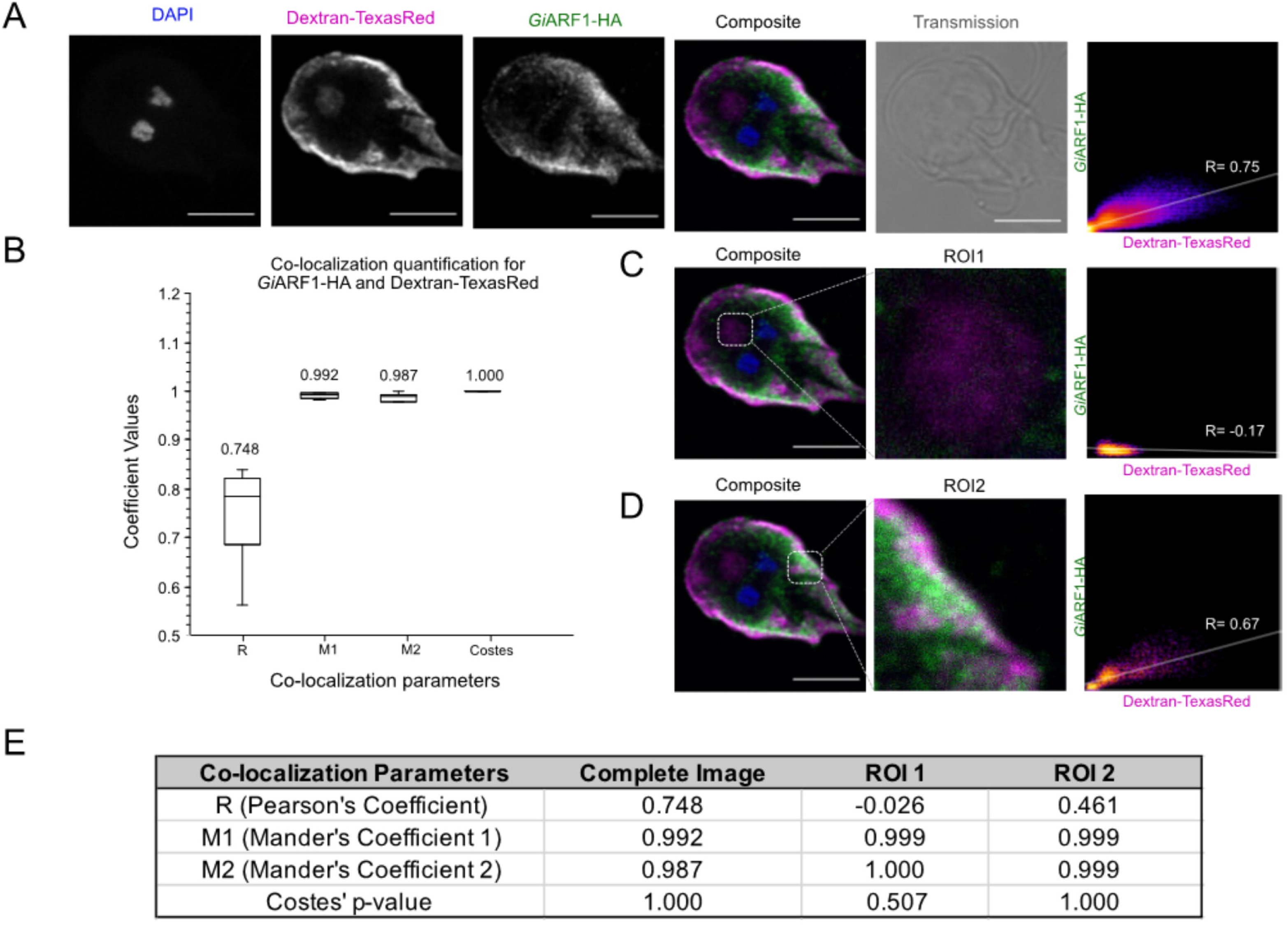
Characterization of *Gi*ARF1-HA subcellular location. (**A**) depicts results from co-labelling immunoprobing experiments with transgenic *Giardia* WB(C6) trophozoites labelled for epitope-tagged *Gi*ARF1-HA (green) and Dextran-TxR (magenta). (**B**) Signal overlap analyses were performed on whole-cell images to determine co-localization between *Gi*ARF1-HA and Dextran-TxR. Boxplot depiction of the calculated R, M1, M2, and Costes’ p-values across ≥10 analyzed cells, along with their mean values indicated on top, is also provided. (**C**) Signal overlap analyses were performed with the bare zone (ROI1) to determine co-localization between *Gi*ARF1-HA and Dextran-TxR. (**D**) Signal overlap analyses were performed at the cortical PV/PECs (ROI2) to determine co-localization between *Gi*ARF1-HA and Dextran-TxR. (**E**) Summary of average co-localization parameters that were calculated for whole-cell image, ROI1, and ROI2 using ≥10 cells. Scale bars: 5 µm. Optical slices were acquired from the middle of the cell for a maximum projection view of the nuclei and the bare zone. All images were acquired using the Leica SP8 x STED laser scanning confocal microscope under the 100x oil immersion objective lens.

### ARF1FA and ARF1FB1 localize to PECs in a pattern similar but not identical to ARF1

All assemblages of *G. intestinalis* were found to possess numerous additional paralogues of ARF1 that arose at the base of the fornicate lineage [12]. Although there were some differences in the number of individual paralogues, ARF1FA and ARF1FB1 were conserved across all fornicates and in *Giardia intestinalis* AWB. These are currently annotated as ARF3 and ARF2, respectively, on the GiardiaDB, but will be referred to as *per* the former nomenclature to reflect shared ancestry with the other fornicate ARF1F proteins. To understand how these proteins localize compared to *Gi*ARF1, localization studies were again performed by generating stably transfected transgenic lines expressing C-terminally HA-tagged variants of these proteins. Overall numbers of cells expressing reporters *Gi*ARF1FA-HA and *Gi*ARF1FB1-HA were similar as for *Gi*ARF1-HA, whereby 89% and 79% of the cells, respectively, had fluorescent signals (Supplementary Table 6; Supplementary Figure 4C-D).

Immunolocalization analyses revealed slightly different patterns of protein localizations for the three ARF paralogs. *Gi*ARF1FA-HA and *Gi*ARF1B1-HA both localized to the PEC regions, but in distinct patterns. Like GiARF1-HA, *Gi*ARF1FA-HA was seen in regions of the cortical PECs but, in contrast, was also detected at the bare zone (Figure 4A).

**Figure 4.**
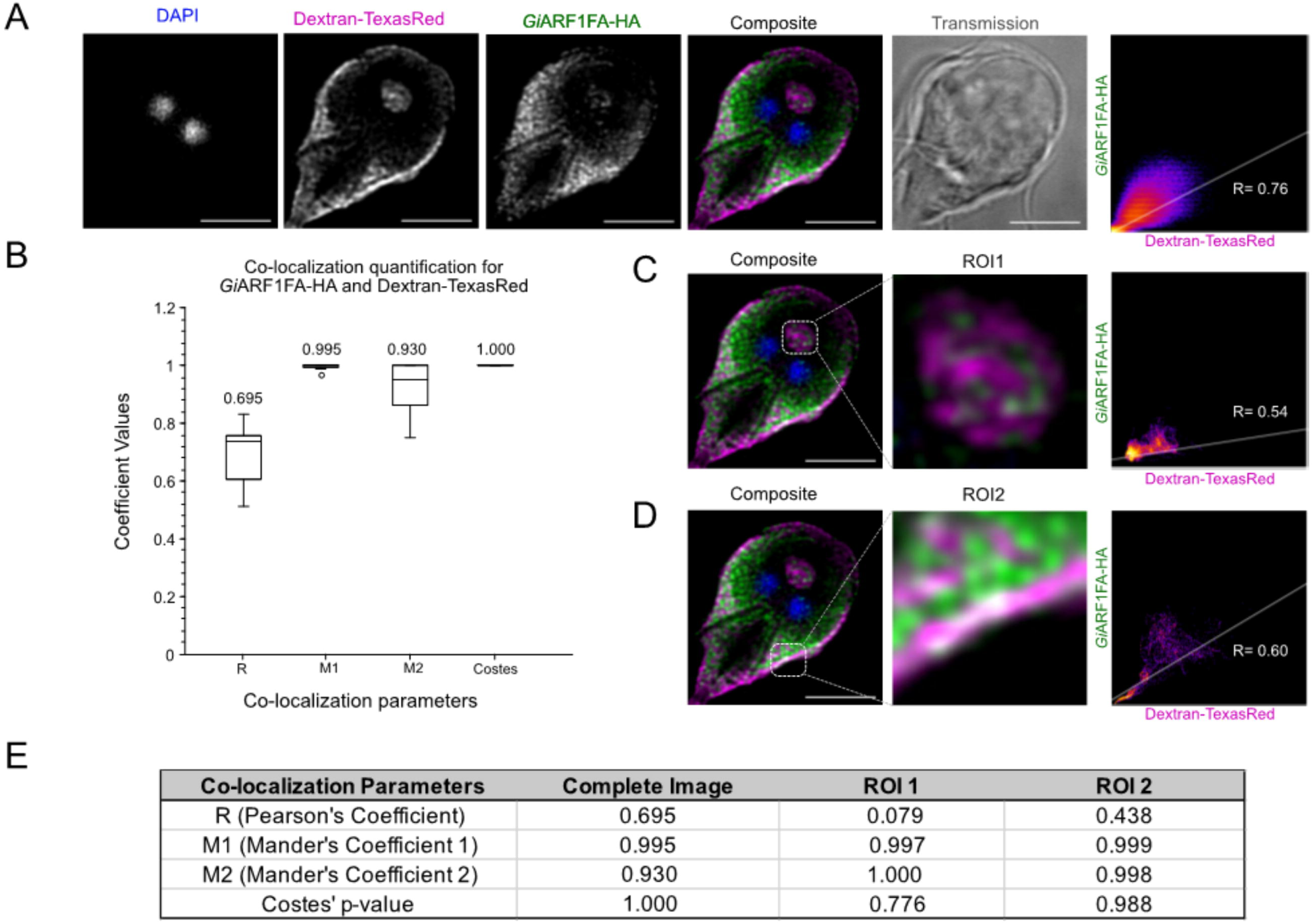
Characterization of *Gi*ARF1FA-HA subcellular location. (**A**) depicts results from co-labelling immunoprobing experiments with transgenic *Giardia* WB(C6) trophozoites labelled for epitope-tagged *Gi*ARF1FA-HA (green) and Dextran-TxR (magenta). (**B**) Signal overlap analyses were performed on complete images to determine co-localization between *Gi*ARF1FA-HA and Dextran-TxR. Boxplot depiction of the calculated R, M1, M2, and Costes’ p-values across ≥10 analyzed cells, along with their mean values indicated on top, is provided. (**C**) Signal overlap analyses were performed at the bare zone (ROI1) to determine co-localization between *Gi*ARF1FA-HA and Dextran-TxR. (**D**) Signal overlap analyses were performed at the cortical PV/PECs (ROI2) to determine co-localization between *Gi*ARF1FA-HA and Dextran-TxR. (**E**) Summary of average co-localization parameters calculated for the whole image, ROI1, and ROI2 using ≥10 cells. Scale bars: 5 µm. Optical slices were acquired from the middle of the cell for a maximum projection view of the nuclei and the bare zone. All images were acquired using the Leica SP8 x STED laser scanning confocal microscope under the 100x oil immersion objective lens.

Quantification of signal overlap in both regions of interest and the whole image was performed with Dextran-TxR and anti-HA co-labelling (Figure 4A). Positive Pearson’s correlation was observed in complete cell image analyses and ranged between 0.51 to 0.83 (average» 0.7; Figure 4A, B, and E). Significant overlap between signals in channels corresponding to *Gi*ARF1FA and Dextran-TxR were confirmed as high M1 and M2 coefficients (M1 average» 0.99 and M2 average» 0.93). All three observations were considered true positives, as the average Costes’ p-value equaled to 1. Unlike *Gi*ARF1-HA, some visible bare zone localization (ROI1) was also observed, albeit not as prominently, as positive Pearson’s coefficients were obtained across at least 10 cells (Supplementary Table 6; Figure 4C and E). On the other hand, a statistically significant association was observed between *Gi*ARF1FA-HA and Dextran-TxR at the cortical PECs (ROI2), as R fell between 0.4 to 0.77 (average» 0.44). Both M1 and M2 ranged between 0.99 and 1 (average» 0.99) with an average Costes’ p-value of 0.99, also suggesting true signal overlap.

Although similar in expression, examination of *Gi*ARF1B1-HA revealed a different pattern of localization to both *Gi*ARF1-HA and *Gi*ARF1FA-HA, in that a diffuse, punctate signal with some PEC overlap was evident (Figure 5). Like the previous two proteins, co-localization was performed with Dextran-TxR to assess bare-zone (ROI1) and cortical PECs region (ROI2) overlap, along with analyses using whole-cell images (Figure 5). Statistical inference from complete image confirms signal overlap between the recombinant ARF1FB1 and Dextran-TxR, as reflected by positive and overall high Pearson’s coefficient values (average» 0.75; Figure 5A, B, and E). Some bare zone-associated signal was also noted, as confirmed by positive R and high M1, M2, and Costes’ p-values (average R» 0.1, M1» 1.00, M2» 1.00, and Costes’ p-value» 0.99; Supplementary Table 6; Figure 5C and E). Finally, as for ARF1FA, a considerable signal overlap was observed at the cortical PECs (Average R» 0.46, M1» 0.99, M2» 0.99 and Costes’ p-value» 0.96; Figure 5D and E).

**Figure 5.**
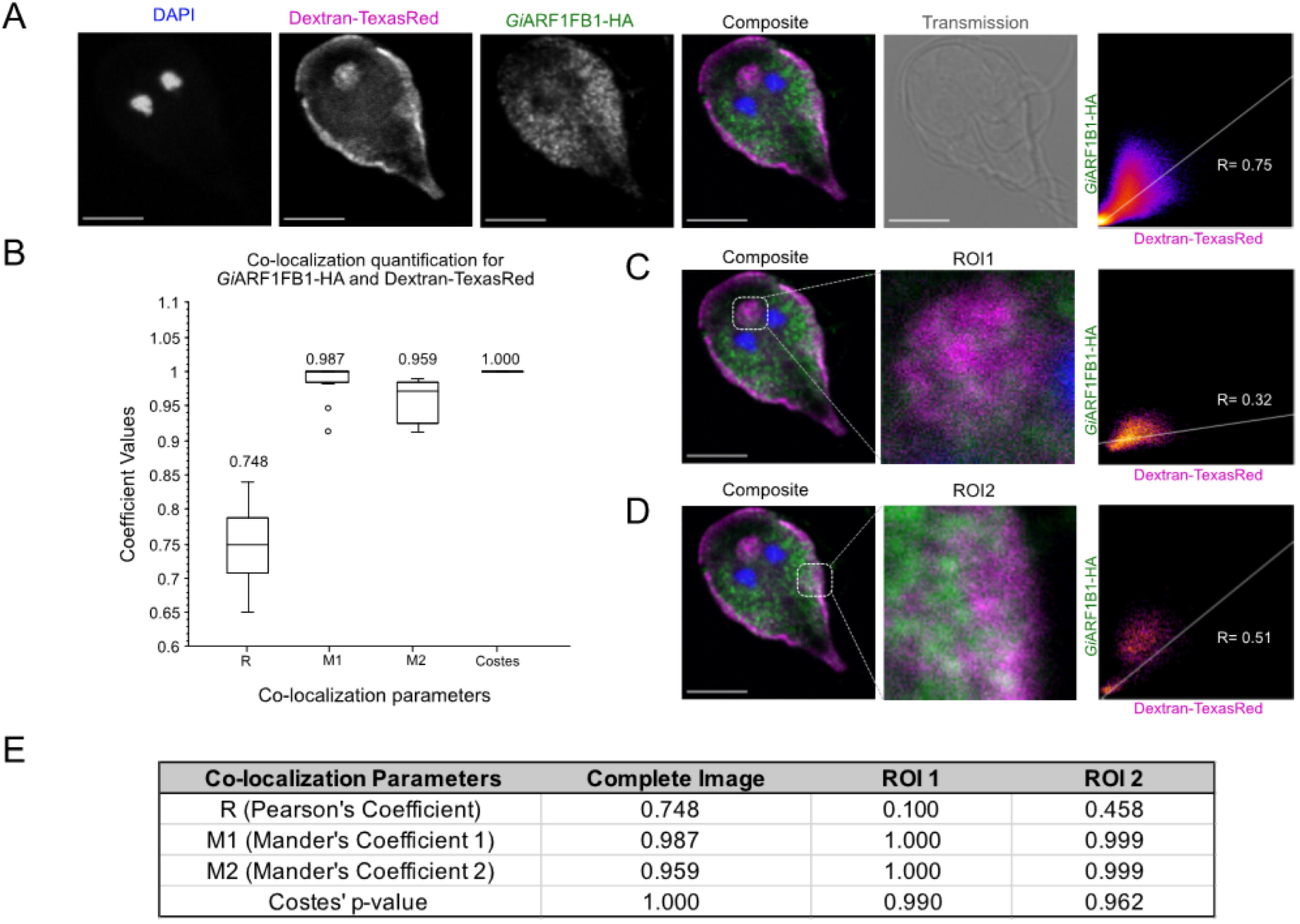
Characterization of *Gi*ARF1FB1-HA subcellular location. (**A**) depicts results from co-labelling immunoprobing experiments with transgenic *Giardia* WB(C6) trophozoites labelled for epitope-tagged *Gi*ARF1FB1-HA (green) and Dextran-TxR (magenta). (**B**) Signal overlap analyses were performed on whole-cell images to determine co-localization between *Gi*ARF1FB1-HA and Dextran-TxR. Boxplot depiction of the calculated R, M1, M2, and Costes’ p-values across ≥10 analyzed cells, along with their mean values indicated on top, is provided. (**C**) Signal overlap analyses were performed at the bare zone (ROI) to determine co-localization between *Gi*ARF1FB1-HA and Dextran-TexasRed. (**D**) Signal overlap analyses were performed at the cortical PV/PECs (ROI2) to determine co-localization between *Gi*ARF1FB1-HA and Dextran-TexasRed. (**E**) Summary of average co-localization parameters were calculated for the whole image, ROI1, and ROI2 using ≥10 cells. Scale bars: 5 µm. Optical slices were acquired from the middle of the cell for a maximum projection view of the nuclei and the bare zone. All images were acquired using the Leica SP8 x STED laser scanning confocal microscope under the 100x oil immersion objective lens.

Taken together, although the three ARF paralogues all appear to localize in close proximity to PECs, the extent to which they do so is not the same. While we note that addition of an epitope tag at the C-terminus and/or the level of expression of the exogenously expressed proteins, particularly compared to those of the endogenous proteins, can result in some alterations to localization, the comparisons of localization differences between the three ARFs are expected to result from differences in their primary sequences.

### GiARFGAP1 and GiCYTHa localize to PECs

As was done for the ARFs, *Gi*ARFGAP1 and *Gi*CYTHa were transiently expressed in Giardia as full-length, recombinant proteins with a C-terminal HA-epitope. Trophozoites expressing *Gi*ARFGAP1-HA were checked for protein expression by staining and counting fixed cells for HA signal, and 79% of cells were found to express the recombinant protein (Supplementary Table 6; Supplementary Figure 4E and G). Similarly, 82% of the trophozoites transfected with the plasmid directing expression of GiCYTHa-HA per positive for HA staining (Supplementary Figure 4F and G).

In mammalian cells, ARFGAP1 regulates ARF1 at the Golgi by interacting with the δ-COP subunit of COPI for retrograde traffic. Given the lack of canonical Golgi bodies in Giardia and the putative connection between ESVs and Golgi, it is possible that ARFGAP1 may act as the ARF GAP necessary for regulating the canonical ARF1 in Giardia’s ESVs during encystation. However, its role in the trophozoite stage is entirely unclear. A detailed inspection of *Gi*ARFGAP1-HA expressing trophozoites revealed a punctate protein distribution at PECs, with some signal dispersal in the cytosol (Figure 6). PEC proximity was evaluated further by co-labeling *Gi*ARFGAP1-HA expressing cells with Dextran-TxR. Whole image co-localization quantification yielded positive Pearson’s correlation values and was accompanied with high M1 and M2 coefficients and Costes’ p-values (average R» 0.51, M1» 1.00, M2» 1.00, and Costes’ p-value» 1), suggesting likely signal overlap between *Gi*ARFGAP1-HA and Dextran-TxR (Supplementary Table 6; Figure 6A and E). Bare zone (ROI1) and cortical PEC (ROI2) associations were assessed through regions of interest analyses. Although strong bare zone association was not observed (average R» −0.03, M1» 1.00, M2» 1.00, Costes’ p-value» 1), some signal overlap was still present at the cortical PEC region (average R» 0.31, M1» 1.00, M2» 1.00, Costes’ p-value» 0.86) (Supplementary Table 6; Figure 6C, D, and E).

**Figure 6.**
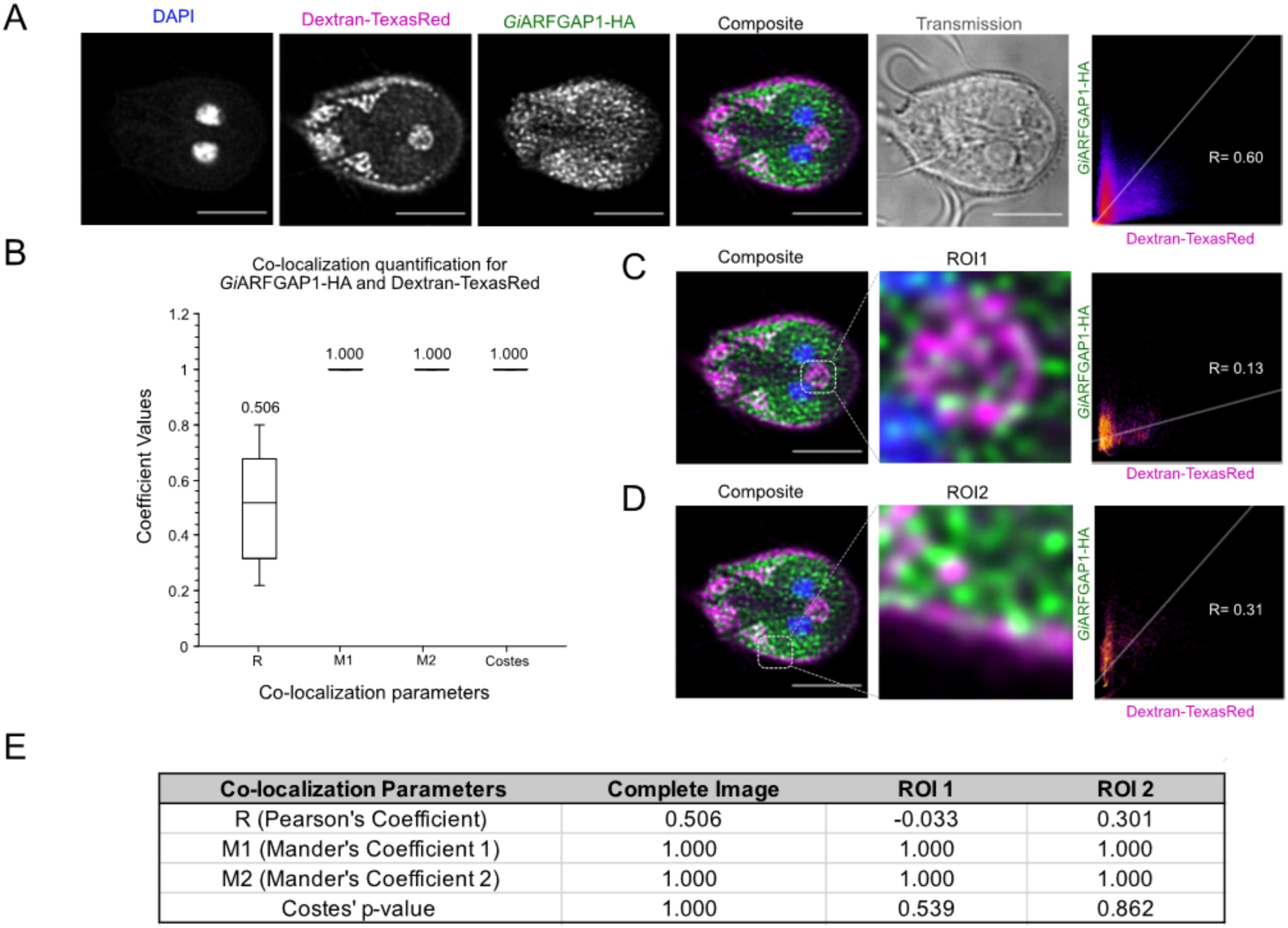
Characterization of *Gi*ARFGAP1-HA subcellular location. (**A**) depicts results from co-labelling immunoprobing experiments with transgenic *Giardia* WB(C6) trophozoites labelled for epitope-tagged *Gi*ARFGAP1-HA (green) and Dextran-TxR (magenta). **(B)** Signal overlap analyses were performed on complete images to determine co-localization between *Gi*ARFGAP1-HA and Dextran-TxR. Boxplot depiction of the calculated R, M1, M2, and Costes’ p-values across ≥10 analyzed cells, along with their mean values indicated on top, is provided. **(C)** Signal overlap analyses were performed with the bare zone (ROI1) to determine co-localization between *Gi*ARFGAP1-HA and Dextran-TxR. **(D)** Signal overlap analyses were performed with the cortical PV/PECs (ROI2) to determine co-localization between *Gi*ARFGAP1-HA and Dextran-TxR. **(E)** Summary of average co-localization parameters were calculated for the whole image, ROI1, and ROI2 using ≥10 cells. Scale bars: 5 µm. Optical slices were acquired from the middle of the cell for a maximum projection view of the nuclei and the bare zone. All images were acquired using the Leica SP8 X STED laser scanning confocal microscope under the 100x oil immersion objective lens.

The pattern of presence/absence of the ArfGEFs varies across Giardia isolates [42], with the commonly used *G. intestinalis* laboratory strain AWB lacking BIG orthologs but possessing two paralogues of Cytohesin (*Gi*CYTHa and *Gi*CYTHb). This has been postulated to suggest varying secretion methods and processes within different Giardia isolates. Intriguingly given the lack of canonical Golgi in Giardia, BIG is needed for traffic within Golgi cisternae and trans-Golgi traffic. By contrast, mammalian Cytohesins activate ARF1 and ARF6 for recruitment onto endosomes by facilitating the GDP to GTP exchange [4,6].

Unlike ARFGAP1, *Gi*CYTHa-HA had an exclusive localization to the PECs. Substantial signal overlap between the recombinant protein and Dextran-TxR was observed both at the bare zone as well as the cortical PEC regions (Figure 7). Quantitative analyses confirmed these qualitative observations where whole-cell image comparisons yielded high positive Pearson’s and Manders’ coefficients, and Costes’ p-values across all sampled cells (average R» 0.68, M1» 1.00, M2» 1.00, and Costes’ p-value» 1; Supplementary Table 6; Figure 7B and E). Examination of the bare zone (ROI1) also showed statistically significant overlap between the two fluorescent signals (average R» 0.40, M1» 1.00, M2» 1.00, and Costes’ p-value» 1; Supplementary Table 6; Figure 7C and E). A similar trend of positive co-localization was also noted at the cortical regions in all sampled trophozoites (average R» 0.56, M1» 1.00, M2» 1.00, and Costes’ p-value» 0.99; Supplementary Table 6, Figure 7D and E).

**Figure 7.**
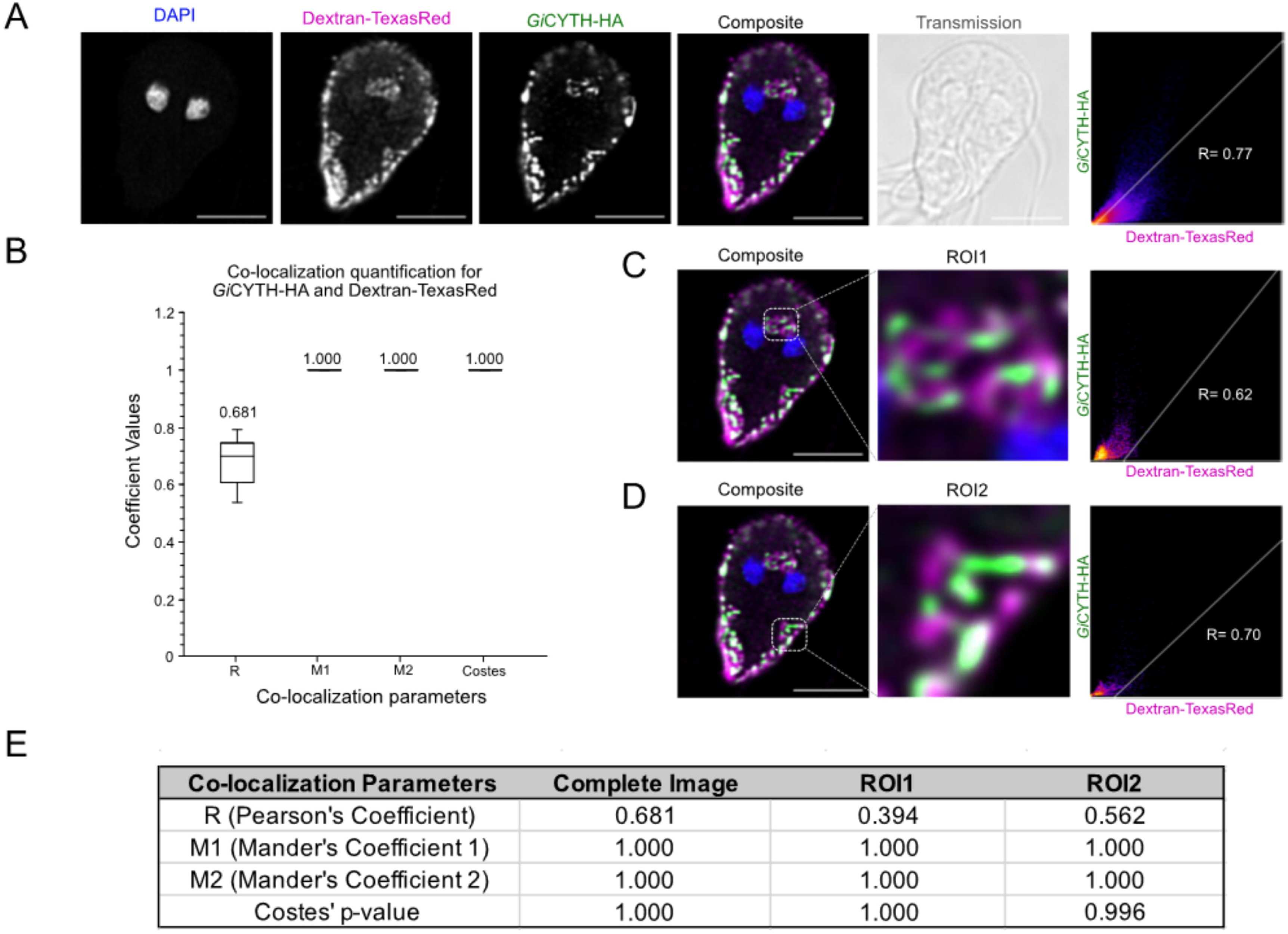
Characterization of *Gi*CYTHa-HA subcellular location. (**A**) depicts results from co-labelling immunoprobing experiments with transgenic *Giardia* WB(C6) trophozoites labelled for epitope-tagged *Gi*CYTHa-HA (green) and Dextran-TxR (magenta). (**B**) Signal overlap analyses were performed on whole-cell images to determine co-localization between *Gi*CYTHa-HA and Dextran-TxR. Boxplot depiction of the calculated R, M1, M2, and Costes’ p-values across ≥10 analyzed cells, along with their mean values indicated on top, is provided. (**C**) Signal overlap analyses were performed with the bare zone (ROI1) to determine co-localization between *Gi*CYTHa-HA and Dextran-TxR. (**D**) Signal overlap analyses were performed at the cortical PV/PECs (ROI2) to determine co-localization between *Gi*CYTHa-HA and Dextran-TxR. (**E**) Summary of average co-localization parameters were calculated for the whole image, ROI1, and ROI2 using ≥10 cells. Scale bars: 5 µm. Optical slices were acquired from the middle of the cell for a maximum projection view of the nuclei and the bare zone. All images were acquired using the Leica SP8 X STED laser scanning confocal microscope under the 100x oil immersion objective lens.

### ARF1-HA associates with Giardia’s vesicle formation machinery at PECs

Collectively, several previously reported *Giardia* vesicle coat proteins, ESCRT components, and now the ARF regulatory proteins demonstrate a considerable degree of overlap in their patterns of cellular localization [15,16,43]. These findings warrant a closer proteomics investigation to define organellar and protein-protein associations. To address this, limited cross-linking co-immunoprecipitation (co-IP) followed by mass spectrometry (MS)-based protein identification experiments were performed, using the HA-tagged variants of the *Giardia* ARF regulatory system proteins as affinity handles, and comparing these data to those from co-IP of non-transgenic cell extracts *i.e.* expressing no HA-tagged protein. Using this approach, the molecular interactions of *Gi*ARF1-HA were first investigated. Specificity of the pulldown was confirmed using immunoblot analyses for both bead and control whole-cell lysate samples. *Gi*ARF1-HA enrichment was validated by the presence of a 22.8 kDa band that stained positive with antibodies to HA, and bait-associating protein conjugates were visualized in the Coomassie-stained gel as more diffuse staining with a higher molecular weight (Figure 8A and B). Beads with bound ARF1-HA and associated proteins were subject to MS, which generated peptide hits that were classified according to overall relative protein abundance to yield (r)iBAQ values (Supplementary table 7), with 195 hits above the 0.01% (r)iBAQ cut-off (Supplementary Table 8). At least 27 of these were well characterized and previously validated PEC-associated proteins [16,17,43–48] (Supplementary Table 9). Most of these PECs-associated proteins belong to families of vesicle formation and fusion machinery. Hetero-tetrameric adaptor complex (HTACs) components were enriched in both replicates, namely COPI components, g-COP, d-COP, a-COP, and b’-Cop, and AP-2 subunits, a2 and m2 (Supplementary Table 9; Figure 8C). Retromer and ESCRT subunits, Vps35 and ESCRTIIIA-Vps31 and ESCRTIIIA-Vps4 [43], respectively, were also present (Supplementary Table 9; Figure 8C). Unexpectedly, in addition to the vesicle formation machinery, several small GTPases that are necessary for catalysis of vesicle fusion dynamics were also identified. Many of these were typically Golgi- and/or endosome-associated; including Rab1, Rab2, and Rab11 (Supplementary Table 9; Figure 8C). The large GTPase, Dynamin, which has been implicated in the PECs-PM membrane fusion dynamics, was also detected in this screen [16,49].

**Figure 8.**
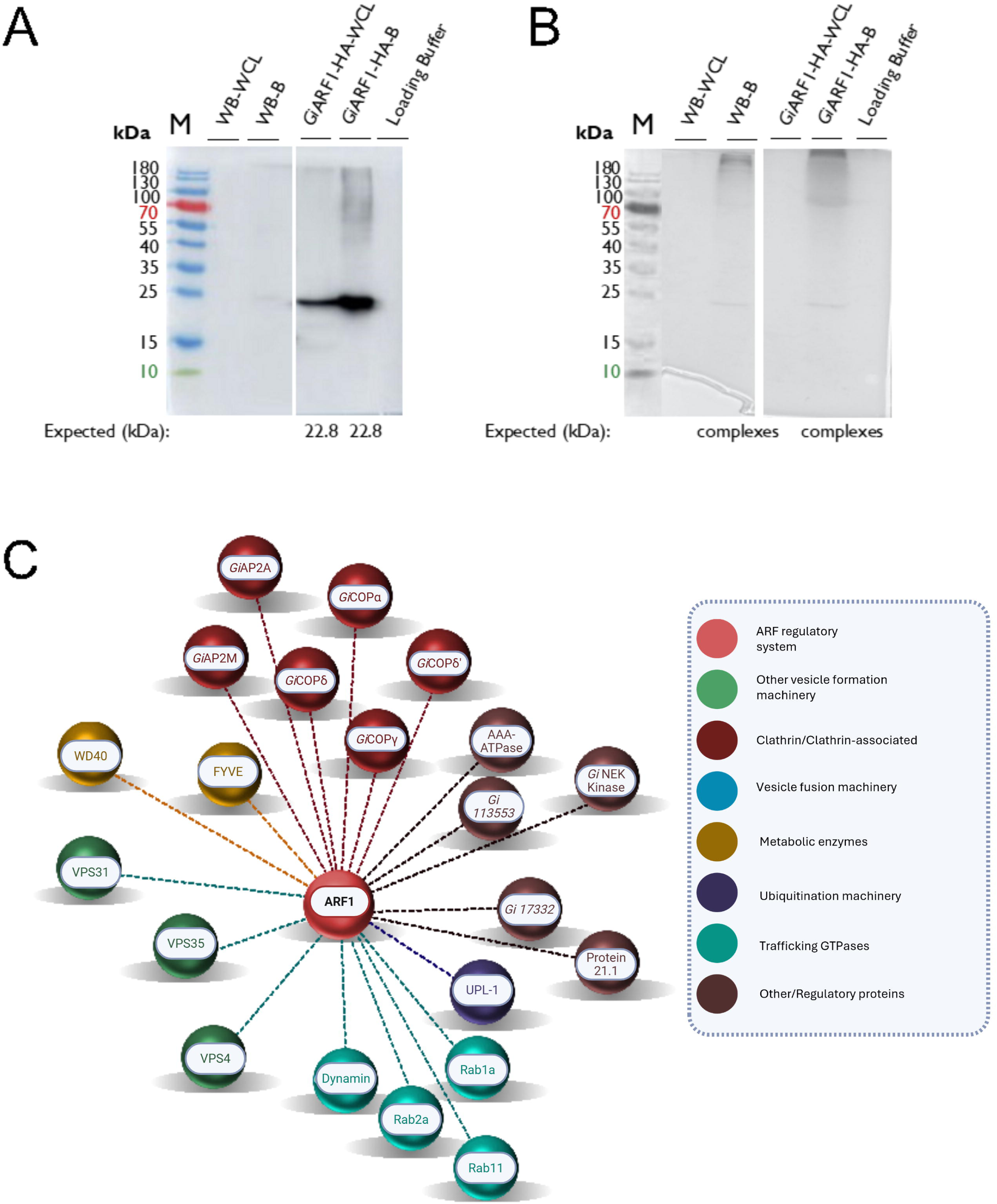
Co-immunoprecipitation and mass spectrometry analyses to determine *Gi*ARF1-HA protein-protein interactions. **(A)** Western blot confirmation of bait pulldown (*Gi*ARF1-HA) in bead and cell lysate samples is depicted, compared to the non-transgenic (WB) control samples, which were performed using SDS-PAGE. An expected band size corresponding to the molecular weight of *Gi*ARF1-HA (22.8 kDa) was observed. **(B)** Coomassie staining was performed to visualize cross-linked protein complexes in bead samples from non-transgenic and transgenic cells. **(C)** The interactome of the experimentally validated PV/PEC-associating proteins enriched in the *Gi*ARF1-HA pulldown and identified by LC/MS is depicted. A threshold of 0.01% (r)iBAQ was used to determine biologically relevant interactions and are indicated by the dashed lines. Abbreviations used: M, Thermo Fisher PageRuler pre-stained protein ladder (10 to 180 kDa), WB, non-transgenic control cells, WCL, whole-cell lysate, B, beads.

Aside from traffic complexes, PEC membrane markers were also present in the co-IPs. Previous investigations characterized numerous phosphoinositide-binding proteins, which are adaptors that bind to specific membrane phospholipids to recruit clathrin and coat proteins for clathrin-mediated endocytosis [17]. In *Giardia*, these PIP-binding proteins are associated with PECs in conjunction with clathrin heavy chain [17]. Two of the previously investigated giardial PIP-binding proteins, FYVE domain-containing protein (GL50803_16653) and WD40 domain-containing protein (GL50803_10822), were also enriched in the *Gi*ARF1-HA interactome [17] (Figure 8C). These findings are consistent with the fluorescence imaging data, all suggesting a physical proximity/association of GiARF1 with PECs.

### ARF1FA-HA and ARF1FB1-HA interactomes are similar to that of GiARF1

Co-IP and LC/MS experiments were also performed with *Gi*ARF1FA-HA and *Gi*ARF1FB1-HA, to assess the molecular interactions common or unique to each paralogue. As previously done, these data were compared to those from co-IP of non-transgenic cell extracts i.e. expressing no HA-tagged protein. *Gi*ARF1FA-HA enrichment was confirmed by immunoblotting, where a 21 kDa band corresponding to the expected protein size was observed in both bead and pellet samples (Figure 9A). Crosslinked protein complexes were visualized through Coomassie staining and analyzed through LC/MS (Figure 9B).

**Figure 9.**
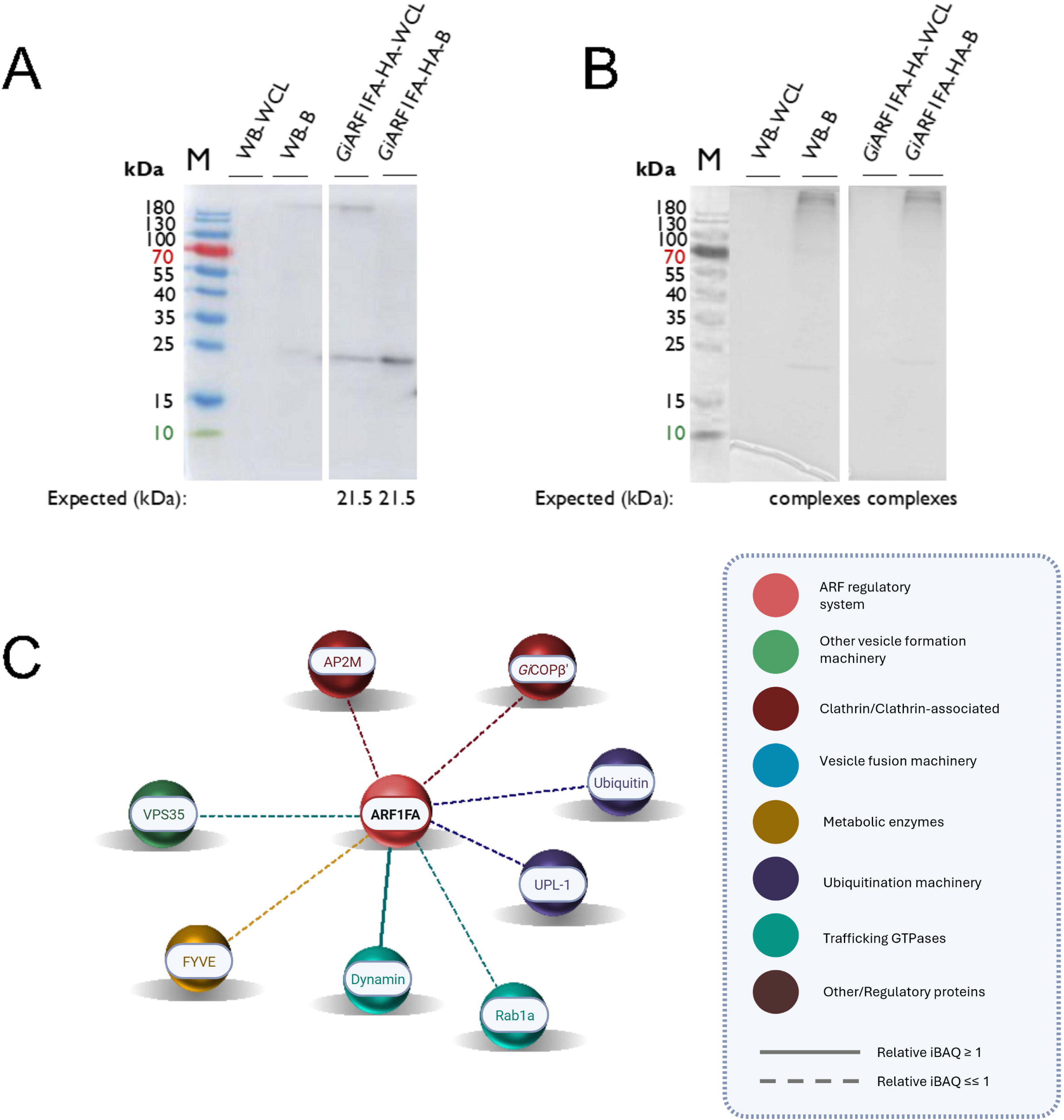
Co-immunoprecipitation and mass spectrometry analyses to determine *Gi*ARF1FA-HA protein-protein interactions. **(A)** Western blot confirmation of bait pulldown (*Gi*ARF1FA-HA) in bead and cell lysate samples is depicted, compared to the non-transgenic (WB) control samples, which were performed using SDS-PAGE. An expected band size corresponding to the molecular weight of *Gi*ARF1FA-HA (21.5 kDa) was observed. **(B)** Coomassie staining was performed to visualize cross-linked protein complexes in bead samples from non-transgenic and transgenic cells. **(C)** The interactome of the experimentally validated PV/PEC-associating proteins enriched in the *Gi*ARF1FA-HA pulldown and identified by LC/MS is depicted. A threshold of 0.01% (r)iBAQ was used to determine biologically relevant interactions and are indicated by the dashed lines. Highly abundant interactions with (r)iBAQ of ≥1% are indicated with solid lines. Abbreviations used: M, Thermo Fisher PageRuler pre-stained protein ladder (10 to 180 kDa), WB, non-transgenic control cells, WCL, whole-cell lysate, B, beads.

A total of 132 proteins were enriched in pulldown studies of GiARF1FA-HA, of which 15 have known PECs functions (Supplementary Tables 4 and 5). Identical to *Gi*ARF1-HA, several of these associations belonged to HTAC components such as AP-2m, β-COP, and retromer-Vps35 (Supplementary Tables 3-5; Figure 9C). Other interaction partners shared by both GiARF1-HA and GiARF1FA-HA included Dynamin, Rab1a, and *Gi*FYVE. Notably, unique to this interactome was the presence of ubiquitin, which is typically necessary for protein post-translational modification and cargo targeting to endo-lysosomal compartments. Overall, although the interactions were not identical, a demonstrable overlap in the types of machinery present in both *Gi*ARF1 and *Gi*ARF1FA datasets existed (Supplementary table 8).

We next performed analogous protein enrichments using *Gi*ARF1FB1-HA co-IP. Immunoblotting revealed a band of 19 kDa, corresponding to the expected size for HA-tagged ARF1FB1, in bead and pellet samples (Figure 10A). Immunoblotting and Coomassie staining also confirmed pulldown of cross-linked complexes (Figure 10B). LC/MS of *Gi*ARF1FB1-HA IPs identified 257 *Giardia* proteins, of which 23 are PEC-associated (Supplementary Tables 3-5). Once again, several PECs-localizing vesicle formation machineries, such as ESCRTIIIA-Vps31 and retromer-Vps26 and Vps35, associated with *Gi*ARF1FB1 (Supplementary Tables 3-5; Figure 10C). However, several important differences were noted, compared to the *Gi*ARF1 and *Gi*ARF1FA datasets. Most notably, none of the HTAC machinery was enriched (Supplementary Tables 3-5; Figure 10C). Unlike the other ARFs, Dynamin and Rab11 were among the strongest interacting partners with >1% (r)iBAQ. Several Rabs were also detected (*i.e.,* Rab1, Rab32, and Rab2a), but also the SAR1 GTPase (typically found at the ER; Figure 10C). Intriguingly, associations with several Q and R SNAREs, namely Syntaxin 1 and Synaptobrevin, which were not identified in the other two ARF datasets, were enriched in our ARF1FB1-HA co-IP dataset (Figure 10C). These proteins typically assemble within the SNARE complex for vesicle fusion dynamics at the plasma membrane for cargo exocytosis [50]. As with *Gi*ARF1 and *Gi*ARF1FA, associations with proteins such as *Gi*FYVE and *Gi*PROP1 shown to bind phosphorylated derivatives of phosphatidylinositol phosphates (PIPs), were also observed [17,51]. However, unique to this co-IP was enrichment of a third PIP-binding protein, GiPXD3 (Figure 10C). Finally, *Gi*ARF1FB1 also associated with ubiquitination machinery (ubiquitin and UPL-1) as well as NEK kinases previously reported at PECs (Figure 10C) [25].

**Figure 10.**
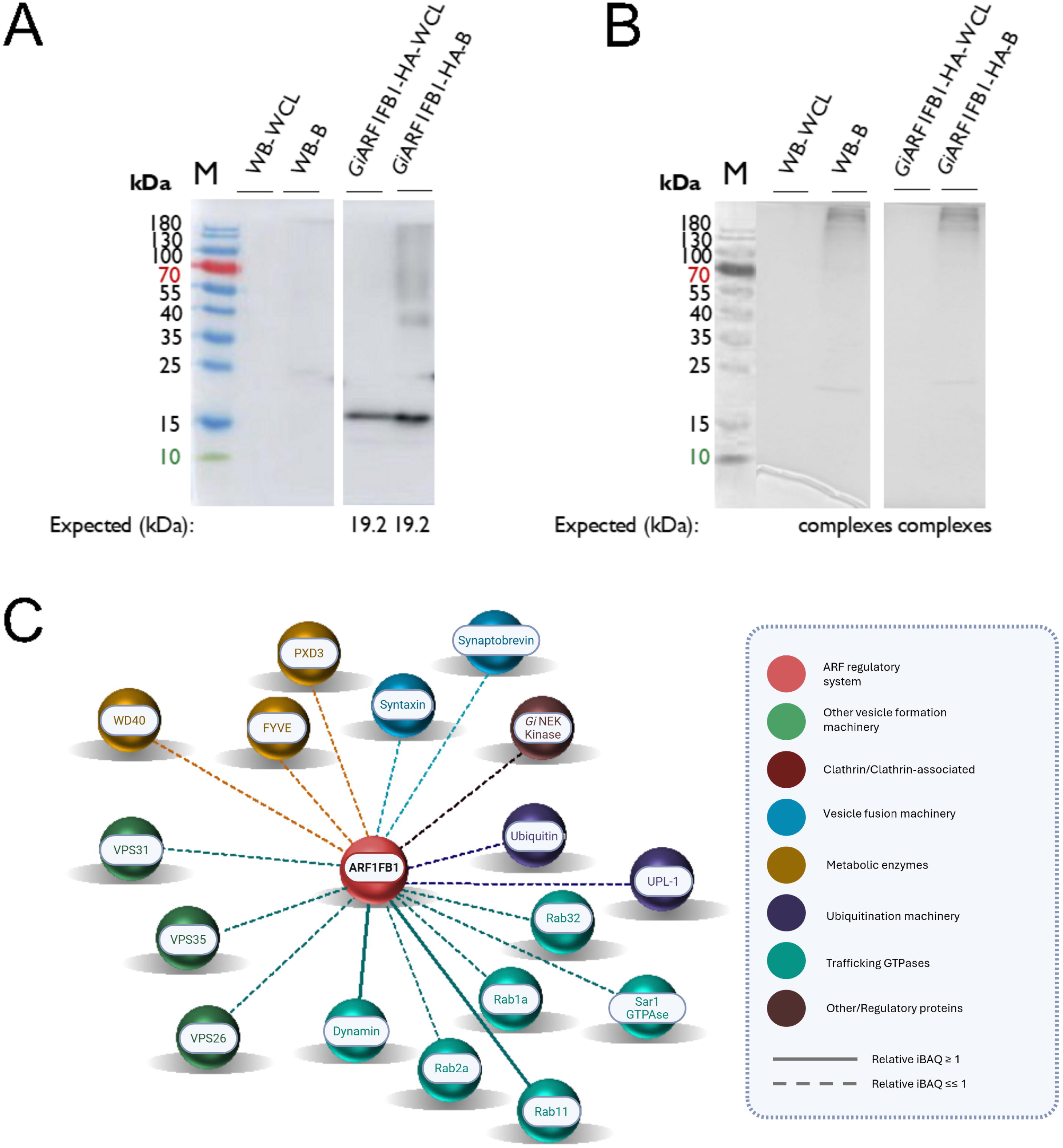
Co-immunoprecipitation and mass spectrometry analyses to determine *Gi*ARF1FB1-HA protein-protein interactions. **(A)** Western blot confirmation of bait pulldown (*Gi*ARF1FB1-HA) in bead and cell lysate samples is depicted, compared to the non-transgenic (WB) control samples, which were performed using SDS-PAGE. An expected band size corresponding to the molecular weight of *Gi*ARF1FB1-HA (19.2 kDa) was observed. **(B)** Coomassie staining was performed to visualize cross-linked protein complexes in bead samples from non-transgenic and transgenic cells. **(C)** The interactome of the experimentally validated PV/PEC-associating proteins enriched in the *Gi*ARF1FB1-HA pulldown and identified by LC/MS is depicted. A threshold of 0.01% (r)iBAQ was used to determine biologically relevant interactions and are indicated by the dashed lines. Highly abundant interactions with (r)iBAQ of ≥1% are indicated with solid lines. Abbreviations used: M, Thermo Fisher PageRuler pre-stained protein ladder (10 to 180 kDa), WB, non-transgenic control cells, WCL, whole-cell lysate, B, beads.

### GiARFGAP1 and GiCYTHa associate with PECs machinery

*Gi*ARFGAP1-HA and *Gi*CYTHa-HA also displayed patterns of localization at the PECs. GEFs and GAPs typically regulate the reversible ARF recruitment to organellar membranes by promoting GDP to GTP exchange and cycling back to the cytosolic pool upon hydrolysis of the GTP. Because both *Gi*ARFGAP1-HA and *Gi*CYTHa-HA localized to similar regions of the PECs as the ARF paralogues, the possibility of ARF cycle regulation by these proteins at these subcellular interfaces was also probed through proteomics.

Transgenic cells expressing *Gi*ARFGAP1-HA were used for co-IP analysis, as previously done, including co-IP of non-transgenic cell extracts as a negative control. Immunoblotting and Coomassie staining confirmed pulldown of the *Gi*ARFGAP1-HA bait (expected size: ∼20 kDa) and complexed proteins (Figure 11A and B). LC/MS identified an overlapping protein interaction pattern with *Gi*ARF1, *Gi*ARF1FA, and *Gi*ARF1FB1, including vesicle coat components, `α`-COP, β’-COP, γ-COP, and AP-2µ (Supplementary Tables 3-5; Figure 11C). ESCRTIIIA-Vps4, three paralogues of ESCRTIIIA-Vps31, and retromer-Vps35 were also enriched. Of the GTPases, Rab11, SAR1, and Dynamin were detected, along with three PIP-binding proteins (GiPXD3, GiPROP1, and *Gi*FYVE). Interestingly, association of *Gi*ARFGAP1 to ARF1FA and another ARF GAP (AGFG) was also detected (Figure 11C). Although ARFGAP1 was not identified in the ARF1FA interactome, ARF1FA was found in the ARFGAP1 pulldowns. This result is consistent with their direct interaction, as predicted by the actions of their mammalian orthologs, but cannot be proven by co-IP alone. Pulldown of PEC-associating proteins is consistent with the localization results at these organelles and with ARFs and ARFGAP1 acting at those sites.

**Figure 11.**
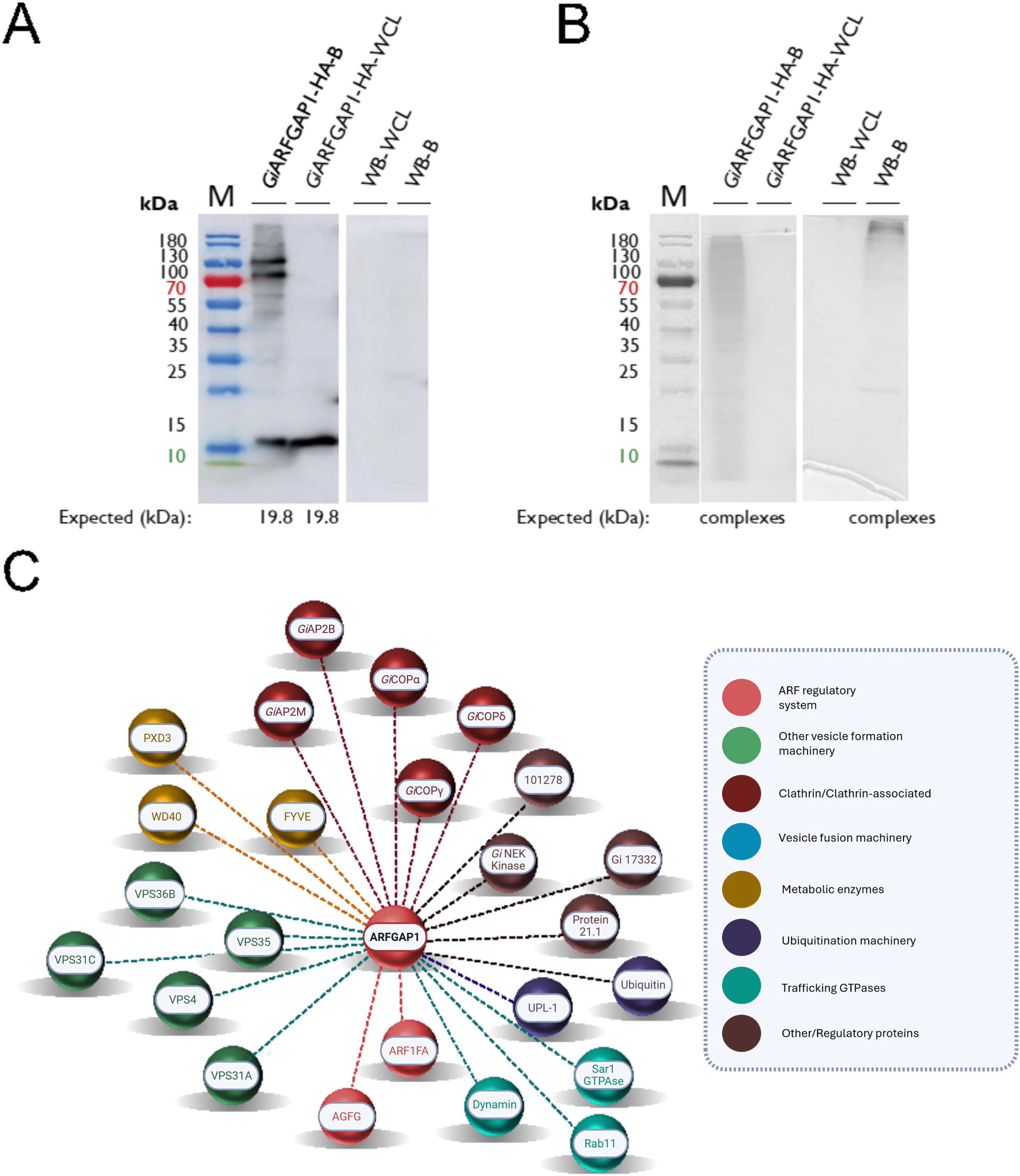
Co-immunoprecipitation and mass spectrometry analyses to determine *Gi*ARFGAP1-HA protein-protein interactions. **(A)** Western blot confirmation of bait pulldown (*Gi*ARFGAP1-HA) in bead and whole-cell lysate samples is depicted, compared to the non-transgenic (WB) control samples, which were performed using SDS-PAGE. An expected band size corresponding to the molecular weight of *Gi*ARFGAP1-HA (19.8 kDa) was observed. **(B)** Coomassie staining was performed to visualize cross-linked protein complexes in bead samples from non-transgenic and transgenic cells. **(C)** The interactome of the experimentally validated PV/PEC-associating proteins enriched in the *Gi*ARFGAP1-HA pulldown and identified by LC/MS is depicted. A threshold of 0.01% (r)iBAQ was used to determine biologically relevant interactions and are indicated by the dashed lines. Abbreviations used: M, Thermo Fisher PageRuler pre-stained protein ladder (10 to 180 kDa), WB, non-transgenic control cells, WCL, whole-cell lysate, B, beads.

*Gi*CYTHa-HA is a much larger protein than the ARFs, with a predicted molecular weight of ∼270 kDa. Instead of observing a single band with a molecular weight of 270 kDa, numerous bands of different sizes were observed (Figure 12A and B) in immunoblots using HA antibodies. Nonetheless, LC/MS confirmed peptides corresponding to *Gi*CYTHa-HA to be highest in enrichment (Supplementary Tables 3-5). Common interactors included 128 proteins shared between the replicate datasets, with a large fraction of these corresponding to ribosomal proteins (Supplementary Tables 3-4). However, others were mostly vesicle traffic proteins, including Rab11, Rab2a, Synaptobrevin, Syntaxin 16, and an unclassified Qa-SNARE. Notably, strong associations with PXD3 and AP-2 were also identified (Figure 12C). Other PEC-associating proteins and transporters were also enriched. Although no ARF or ARF GAP interactions were identified in this screen, these data are consistent with immunofluorescence observations for *Gi*CYTHa-HA at PECs.

**Figure 12.**
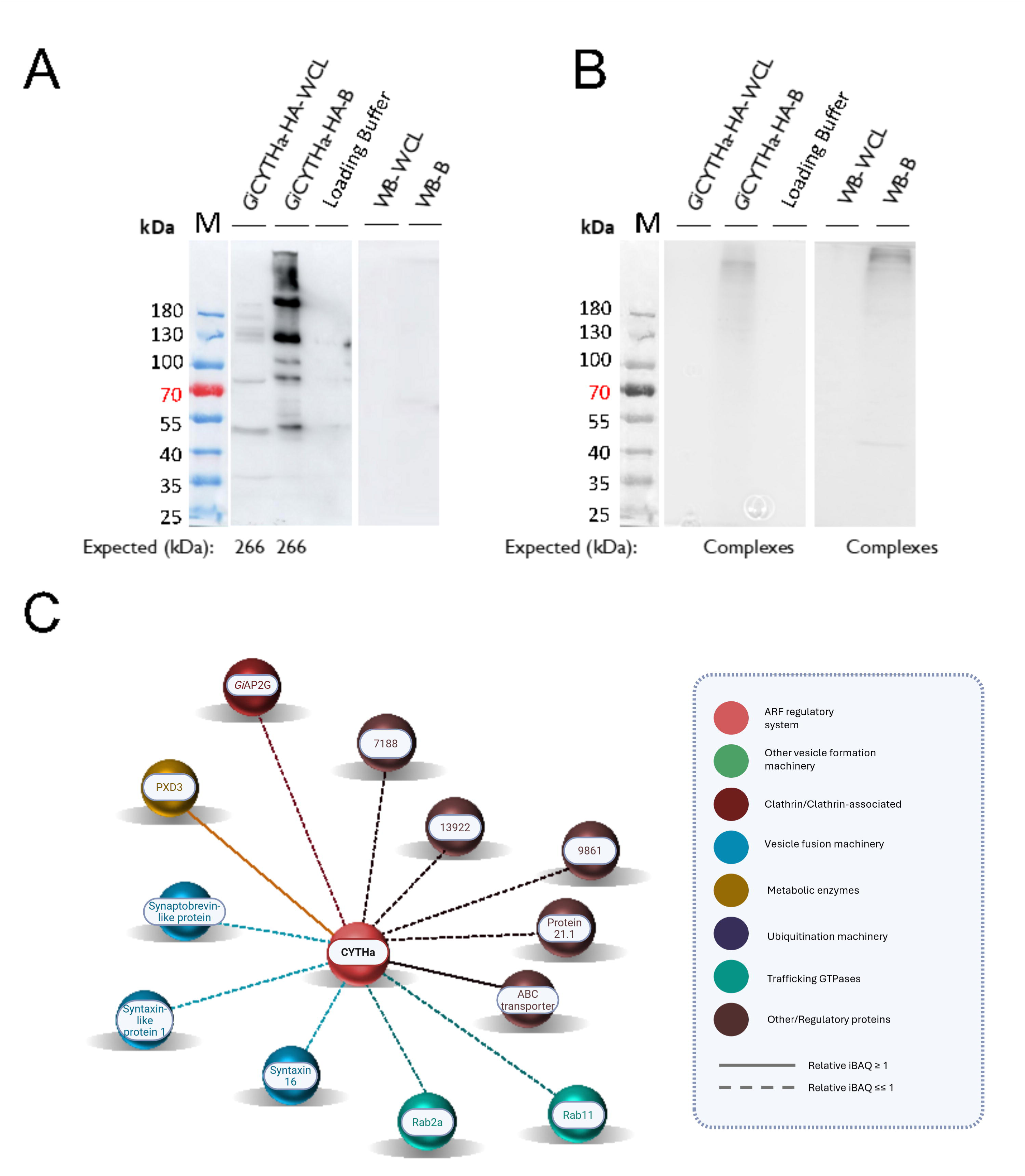
Co-immunoprecipitation and mass spectrometry analyses to determine *Gi*CYTHa-HA protein-protein interactions. **(A)** Western blot confirmation of bait pulldown (*Gi*CYTHa-HA) in bead and whole-cell lysate samples is depicted, compared to the non-transgenic (WB) control samples, which were performed using SDS-PAGE. An expected band size corresponding to the molecular weight of *Gi*CYTHa-HA (226 kDa) was not observed. Instead, numerous bands of variable sizes in both the bead and cell lysate samples were present in the bait-associated lanes. **(B)** Coomassie staining was performed to visualize cross-linked protein complexes in non-transgenic and transgenic bead samples. **(C)** The interactome of the core PV/PEC-associating proteins enriched in the *Gi*CYTHa-HA pulldown and identified by LC/MS is depicted. A threshold of 0.01% (r)iBAQ was used to determine biologically relevant interactions and are indicated by the dashed lines. Highly abundant interactions with (r)iBAQ of ≥1% are indicated with solid lines. Abbreviations used: M, Thermo Fisher PageRuler pre-stained protein ladder (10 to 180 kDa), WB, non-transgenic control cells, WCL, whole-cell lysate, B, beads.

Co-enrichment with each of the five *Giardia* ARF-regulatory system components described above provides supportive evidence for their common association with PECs. To consolidate these findings and determine overlapping networks between these PEC associations, all five proteomics datasets were compared (Supplementary Table 9). Interactions are mainly with traffic proteins that typically mark early and late endosomal pathways. These were AP-2 and COPI components, ubiquitination machinery, ESCRTs, and retromer, with vesicle fusion proteins such as SNAREs and Rabs also identified.

## Discussion

Previous characterizations of *Gi*ARF1 function were limited to its role during encystation [52]. In this study we sought to compare the cellular locations and interactomes of the three ARF1 orthologs in G. intestinalis to allow potential comparisons between this organism with minimized membrane traffic compartmentalization and the well-studied mammalian models. We extended our studies to include an ARF GEF and ARF GAP as the most likely representative of each to be acting with the ARFs. To do this, we used a combination of immunofluorescence microscopy and LC/MS proteomics to define the molecular localization and potential interaction partners of all three ARF paralogues and representative regulatory proteins in vegetative Giardia cells. Taken together, our findings confirm two attributes of the ARF pathway that are conserved in other systems. The first is that the Giardia ARF regulatory system likely modulates the assembly dynamics of vesicle coat complexes around the PECs as the singular endo-lysosomal compartment. The other is that the three GiARF paralogs are predicted to play distinct but likely overlapping roles at PECs, consistent with their overlap in localization and shared co-IPing proteins.

### Conventional roles of the Giardia ARF regulatory system and PEC functional diversification

Although Giardia ARFs and selected regulators localize primarily to PECs, they do so in variable patterns, supporting the hypothesis that different regions of PECs have specialized and distinct functions [15]. Like the early and late endosomal stages in mammalian cells, perhaps characterized by changes in the presence of different RAB proteins (REF), a mechanism of organelle maturation may exist within the static-state PECs. Populations closer to the ER may have *trans*-Golgi network-like functions for the extracellular secretion of the newly synthesized material. Those closer to the plasma membrane, including the bare zone, may be involved in selective uptake and sorting of host macromolecules for retrograde traffic leading to downstream metabolic breakdown. Support for such a model is provided by super-resolution optical microscopy (STORM) and focused ion beam scanning electron microscopy (FIB-SEM) investigations of PECs [15] which identify up to three distinct heterogeneous subpopulations of PECs that vary in both size/volume and shape [15].

Mechanisms of PECs biogenesis and the donor membranes which give rise to them remain a mystery. It is unclear whether these endo-lysosomal organelles originate from the plasma membrane, the ER, or both. The ARF regulatory system may be integral to vesicle-budding and maturation-like processes for PECs neogenesis from either of these compartments. Based upon studies in other model systems it appears likely that the different *Gi*ARF1 paralogues and their regulators may coordinate coat protein assembly and nucleation (*e.g.,* COPII, COPI, adaptins, and retromer), followed by tethering and fusion in a SNARE-dependent manner (*i.e.,* Syntaxins and Synaptobrevins) for synergistic vesicle scission and fusion to yield different PECs-populations. Association with the giardial COPII machinery (*e.g.,* SAR1) and *Gi*Syntaxin 1 and *Gi*Syntaxin16 is a scenario reminiscent of mammalian ARF1-dependent formation of COPII-coated endosomes that arise at ER/ERGIC interfaces for fusion with the *cis*-Golgi in coordination with SNARE-helical bundles [53]. *Giardia* trophozoites that lack stacked-Golgi or ESVs may have stages of PECs, which are functionally analogous to COPII-vesicles or the *cis*-Golgi. Fluorescence microscopy with *Gi*ARF1FB1 supports this notion as the localization of this protein extends close to regions spanning the *Giardia* ER.

*Gi*ARF1 and *Gi*ARF1FA interact with post-Golgi endosomal machinery, which suggests a link between cortical PECs populations and early/late endosomes. Both proteins are associated with components of COPI, AP-2, retromer, and the ESCRT subcomplexes, which typically facilitate the formation and maturation of early/recycling endosomes or multivesicular bodies. Although the role of *Giardia* clathrin has diverged from clathrin-coated vesicle formation and the parasite is devoid of clathrin uncoating factors, vesicle coats are identified at the cortical and bare zone PECs populations near the plasma membrane [16,43,47,54]. Investigations of similar scope have shown PEC populations to be marked and modulated by PIP-binding proteins carrying FYVE, PX, and NECAP1 domains [16,17,55,56], shown to interact with adaptin and ARF-regulatory components in *Giardia* [17].

Several components of the *Giardia* ARF regulatory system are also associated with PIP-binding domains and kinases. Investigations by Cernikova and colleagues demonstrated defects and aberrations in PECs morphology upon depletion of canonical endosomal phospholipids (*i.e.,* PI(3)P, PI(3,4)P2, and PI(3,4,5)P3) which impaired fluid-phase uptake dynamics [17]. In *Dictyostelium*, arrestins with FYVE-domain architecture were implicated in the selective recruitment of lineage-specific ARFA for early endosomal transport of arrestins via recognition and binding of PI(3)P at the plasma membrane [57].

Similarly, in mammalian systems, ARF6 is essential for the activation of PtdIns4P 5′-kinase (PIP5K) and to control cargo sorting of PX domain-containing phospholipase D (PLD) through AP-2 mediated clathrin-dependent endocytosis at the plasma membrane [58]. The findings from this study suggest that *Gi*ARF1 and *Gi*ARF1A have roles in endosome-like traffic processes that occur in synchrony with AP-2, PIP-binding proteins, and kinases.

Finally, all three *Gi*ARFs as well as *Gi*ARFGAP1 interact with the parasite’s retromer components (*i.e., Gi*Vps35 and *Gi*Vps26), reflecting another conventional paradigm of ARF-mediated endosomal traffic to exist in this parasite (Supplementary table 9). In mice fibroblast and HeLa cells, ARF6 regulates recruitment of retromer-mediated traffic of mannose-6-receptors, where ARF6 deletion mutants exhibit aberrant localization of retromer components and cargo away from the TGN pathway, which in turn results in an aberrant tubular morphology of the endosomes [59].

### A connection between the ARF complement and Rab GTPases in Giardia

Our proteomics investigations consistently identified *Giardia* ARFs and their regulators to co-IP with traffic-associated Rabs. This suggests possible signaling-cascade recruitment of Rabs (or *vice versa*) and crosstalk between the two pathways, as first identified in the yeast *S. cerevisiae* [60]. In line with this hypothesis, mammalian ARF4 requires associations with the ARF GAP, ASAP1 and Rab11 at the TGN, which results in cascade recruitment of other Rabs, and ARF- and Rab-interacting effectors (*e.g.,* Rab8 and FIP3) to provide specificity for ciliary traffic of rhodopsin [61]. Emerging lines of evidence in parasite endomembrane systems have also implicated a role of ARF-Rab cross talk. In *Plasmodium*, late endosomal Rab5 and Golgi-associated Rab1 regulate traffic of N-myristoylated adenylate kinase 2 (*Pf*AK2) and transmembrane protein Rifin to the parasite-induced parasitophorous vacuole [62]. This process relies on cargo sorting by *Pf*ARF1 at the parasite ER-exit sites, followed by recruitment of Rab5 and Rab1. Other glimpses of crosstalk between these pathways are evident throughout the eukaryotic tree of life in the form of domain fusions [9,63].

Finally, ARF GAPs and their roles as effectors within the ARF-Rab cascade are also not unprecedented. The first indication that ARF GAPs serve effector functions came from genetic studies in *S. cerevisiae*, when it was found that high copy suppression of hypomorphic ARF mutations was consistently found for the yeast ARF GAPs [64]. Later, ASAP1 was suggested to have indirect effector functions towards GTP-bound ARF6 for downstream GAP activity towards mammalian ARF1 and ARF5 for traffic of paxillin to invadopodia produced by breast tumor cells for their migration and systemic invasion [6]. Numerous studies continue to highlight the effector functions of ARF GAPs towards both ARFs and Rabs [5,7,8]. Therefore, it is not unlikely that similar regulatory mechanisms exist within *Giardia* ARF/Rab pathways, and results from this investigation are likely providing a glimpse into much more convoluted cargo traffic regulation at play within this simplified parasitic cell.

### Significance of paralogous expansions within fornicate ARF regulatory systems

Duplications within the ARF1 GTPase in Fornicata to yield lineage-specific paralogues occurred in conjunction with loss of ARF6 and streamlining within the ARF GEFs and the ARF GAPs [12]. Notably, a similar evolutionary trend of expansion and loss exists across the tree of eukaryotes. For example, in Holozoa, new families of ARFs (*i.e.,* class I, II, and III ARFs), ARF GAPs (*i.e.,* AGAP, ARAP, and GIT), and ARF GEFs (*i.e.,* FBX8, EFA6, and Cytohesins 1-4) arose in parallel to cellular specialization with distinct functions that depend on these different ARF, ARF GAP, or ARF GEF activities [9,11,65].

Although gradual simplification in traffic organelles persists across all fornicates (*i.e.,* loss of stacked Golgi), the presence of endosome-like compartments in most lineages may be indicative of latent ARF-dependent vesicle formation. In *Giardia*, PECs are unique endo-lysosomal organelles whose homology and dynamics underpinning organellar function and neogenesis are still an active area of investigation. This study identifies an additional layer of complexity and identities of the traffic machinery that may be participating in PEC regulation. Identification of the associating molecular players has also cast light onto notions of crosstalk between different GTPase pathways and the involvement of the vesicle fusion machinery. Especially in this latter case, Giardia could therefore serve as a model for a stripped-down system to probe questions that would otherwise be more challenging to address in systems with greater cellular complexity.

## Materials and Methods

### Dataset acquisition of new Metamonada genomes

Updated genome assemblies for *Anaeramoeba flamelloides* and *Anaeramoeba ignava,* as well as transcriptomes for *Barthelona sp. PAP020, Barthelona sp. PCE, Skoliomonas litria, Skoliomonas sp. GEMRC*, and *Skoliomonas sp RCL.* were obtained for all ARF regulatory system searches. Transcriptomes were additionally translated using *ab initio* gene prediction program, GeneMarkS-T under default parameters [66]. Database sources and references from which all genomes and transcriptomes were obtained are summarized in Supplementary Table 4.

### Comparative genomics of ARF1, ARF6, ARF GAPs, and ARF GEFs in new Metamonads

Published query ARF1 sequences were obtained from previously published reports [10,67]: ARF6 (*Homo sapiens*) from the National Center for Biotechnology Information (NCBI), ARF GAP sequences [11] and ARF GEF Sec7 sequences [12]. Homology and HMM searches were performed using the automated comparative genomics software AMOEBAE [68]. Modifications to the default Snakefile include increased maximum hits to sum from 5 to 400, maximum e-value increased from 0.0005 to 0.05, and removal of aasubseq flag to cause processing of entire amino acid sequences. Searches were performed against genomes and transcriptomes described in Supplementary Table 4. AMOEBAE results are summarized in Supplementary Table 2. ARFGAP and ARFGEF positive hits were subjected to domain prediction using the web interface of InterProScan [69].

### Phylogenetic analyses

Identified *Anaeramoeba flamelloides* and *Anaeramoeba ignava, Barthelona sp. PAP020, Barthelona sp. PCE, Skoliomonas litria, Skoliomonas sp. GEMRC*, and *Skoliomonas sp RCL* ARF1 and ARF6 sequences were used to run phylogenetic analysis using a ARF/ARL backbone [30]. Alignments were conducted in MAFFT [70], using the E-INS-I model. Maximum likelihood analyses were performed using nonparametric and ultrafast bootstrapping (1000), as well as Shimodaira–Hasegawa approximate Likelihood Ratio Test (SH-aLRT) [71] (1000) using IQTree v2.3.6 [72]. Bayesian analyses were conducted via MrBayes v3.2.7a [73] with 4 chains, an MCMC generation of 1000000, a sample, print, and diagnostic frequency of 1000, saved branch lengths, and burn in of 150. Multiple runs were conducted during this process of attempting to phylogenetically differentiate ARF1 and ARF6 in these lineages. Preliminary trees available in Supplementary Figures 1-3. To confirm that the ARF1 hits were true positives, manual BLASTp was conducted on the BaSk sequences [74]. Final runs were conducted with this reduced dataset. Shared SH-aLRT and ultrafast bootstrap values from IQTree, and posterior probability values derived from MrBayes were combined and overlayed onto the IQTree topology. IQTree and MrBayes analyses were conducted on the SciComp server maintained by the University of Alberta. Trees were visualized in FigTree v1.4.4 [75], and nod value annotations were prepared in Affinity Designer.

### Giardia trophozoite culture and transfection

Cell culture and transfection of *Giardia intestinalis* strain A, isolate WB clone C6 (ATCC 50803) trophozoites were performed as per standard protocol [16,17,43,76]. Briefly, trophozoites were axenically cultured in Nunc^TM^ polystyrene culture tubes (Thermo Fisher Scientific) containing *ca.* 11 mL of Diamond’s TYI-S-33 medium supplemented with 10% Seraglob bovine serum (Bioswisstec AG), 0.52 mg/mL bovine bile (Sigma B-8381), 10,000 units/mL penicillin-streptomycin (Thermo Fisher), and 22.8 mg/mL ammonium ferric citrate (Fluka 09714; Sigma), and incubated at 37°C until confluent with subculturing performed every 2-4 days (Keister, 1983). 1×10^6^ mid-log phase vegetative trophozoites were harvested by placing culture tubes on ice for 30 to 45 minutes, followed by gentle hitting and inversion to dislodge the adherent cells [77]. Parasite transgenic lines were prepared by first collecting non-transgenic trophozoites cells by centrifugation at 900 *x g* at 4°C for 10 minutes, followed by electroporation (350V, 960 µF, 800Ω) of 15 μg of pPacV-Integ-based circular plasmid vectors (episomes) expressing reporter constructs that were prepared in electrocompetent *Escherichia coli* D10HB (Thermo Fisher Scientific). Transgenic parasites were selected and sub-cultured with the addition of 40 μL Puromycin (50 μg/ml; InvivoGen) in the culture medium. Parasites were harvested 1.5 to 2 weeks post-transfection and tested for reporter construct expression through immunofluorescence assays and widefield fluorescence microscopy using Leica DM5500 B microscope under 100X oil immersion lens (Leica Microsystems).

### Genomic DNA isolation and cloning for episomal vector construct generation

Genomic DNA was isolated from non-transgenic trophozoites by first harvesting confluent trophozoite cultures, as detailed above. Cells were collected as pellets and resuspended in a buffer solution containing 50 nM Tris (pH 8), 10 mM Ethylenediaminetetraacetic acid (EDTA), and 1.5 μL of 100 mg/mL RNase, and vortexed until a homogenous slurry was observed. 200 mM NaOH and 1% SDS were added to this solution and incubated at room temperature for two minutes, then resuspended in 3M potassium acetate (pH 5) followed by centrifugation at 14000 *x g* for 15 minutes at 4°C. gDNA was precipitated out of solution by adding 900 μL of 70% cold ethanol, after which supernatant was discarded, and the resulting DNA pellet was dissolved in 20 μL of ddH_2_0. Isolated gDNA yield and purity were measured using a Nanodrop spectrophotometer (ThermoFisher). Ten nanograms of the isolated gDNA were used for polymerase chain reaction (PCR) amplification of the open reading frames belonging to *Gi*ARF1 (GL50803_7789), *Gi*ARF1FA (GL50803_13930), *Gi*ARF1FB1 (GL50803_75620), *Gi*CYTHa (GL50803_17192), and *Gi*ARFGAP1 (GL50803_2834), along with their native promoters and a C-terminal hemagglutinin tag (HA). As discussed earlier, the C-terminal tagging strategy was employed as the ARF N-terminus is necessary for N-myristoylation and membrane recruitment. PCR amplification was performed using designated oligonucleotide pairs as listed in Supplementary Table 5, synthesized by Microsynth (Balgach, Switzerland). Due to the large predicted open reading frame of *Gi*CYTHa (GL50803_17192), a cloning strategy using the Gibson Cloning Kit Protocol (Addgene) was employed. For all ORFs, promoter sequences were derived from 150 to 200 base pairs upstream of the predicted start codon. The resulting inserts were cloned into a pPacV-Integ modified vector containing XbaI and PacI restriction enzyme recognition sites, the giardial glutamate dehydrogenase promoter, and a puromycin resistance cassette for constitutive episomal expression of each *Giardia* ARF regulatory system protein and antibiotic selection of transgenic cells [16]. Detailed vector maps for all constructs are provided in supplementary Figure 6 while construct sequences are provided in supplementary files 3-7.

### Indirect immunofluorescence and fluid-phase uptake assays

Transgenic lines and non-transgenic control cells were cultured in 1 x Nunc^TM^ T-25 polystyrene flasks (Thermo Fisher) per line until parasites reached confluency. Cells were harvested by cooling and detaching on ice for 30 minutes, followed by centrifugation at 900 *x g* for 10 minutes. The resulting cell pellets were washed with cold PBS and fixed with 3% formaldehyde (Sigma) in phosphate-buffered saline (PBS) for 1 hour, followed by quenching using 0.1M Glycine in PBS for 5 minutes. Fixation and quenching were both performed at room temperature. Cells were permeabilized using 2% bovine serum albumin (BSA)/0.2% Triton X-100 in PBS for 20 minutes at room temperature and blocked in 2% BSA in PBS for two hours. Antibody incubations were performed using a primary rat-derived monoclonal anti-HA high-affinity antibody (dilution 1:250; Roche) and a secondary goat-derived anti-Rat IgG (H+L) conjugated to Alexa Fluor 488 (AF488) (dilution 1:250; Thermo Fisher). Both antibody solutions were prepared in 2% BSA/0.2% Triton X-100 in PBS, and incubations for each was performed for one and a half hour at room temperature on a rotating shaker in the dark. Between each antibody incubation, washes were performed using 1% BSA/0.1% Triton X-100 in PBS, after which samples were fixed in 10 to 40 μL Vectashield (Reactolab) containing 4′-6-diamidino-2-phenylindole (DAPI) for nuclear staining and cell suspension. Three microlitres of the cell suspension were aliquoted onto microscopy glass slides and covered with 22 mm x 22 mm coverslips which were sealed with nail varnish.

PEC labelling was performed using the fluid-phase marker Dextran (10,000 MW) conjugated to Texas Red (Cat. No. D1863, Thermo Fisher), as previously described [49]. Briefly, confluent T-25 flasks of transgenic and non-transgenic trophozoites were cultured and harvested as described above. Cell pellets were resuspended in *ca.* 100 μL of freshly supplemented TYI-S-33 medium to which Dextran-Texas Red (Dextran-TxR) was added to a final concentration of 2 mg/mL. Incubations were performed at 37°C for 30 minutes to allow for the uptake of extracellular Dextran-TxR into the PECs lumen, which was then halted by placing the cells on ice for 15 minutes and washing with cold PBS. Cells were chemically fixed with 3% formaldehyde in PBS followed by permeabilization and immunoprobing with primary and secondary antibodies, as per standard immunofluorescence assay protocol described above.

### Laser Scanning Confocal Microscopy and image analysis

Visualization of recombinant proteins probed with primary and secondary antibodies was performed using a Leica SP8 x STED super-resolution microscope configured with white light lasers (excitation wavelength between 470 nm to 670 nm), photomultiplier tubes, and HyD detectors, under HC PL APO 100X/1.44 oil immersion objective lens (Leica Microsystems). Appropriate excitation and emission settings were used for the gating and visualization of green (Alexa Fluor 488), red (Texas Red), and blue (DAPI) channels. Brightfield images were acquired using the transmission/differential inference contrast settings (PMT Trans). Laser pinhole opening size was set to Airy 1, and images were acquired with line averaging set to 4 to optimize the signal-to-noise ratio. More than 100 cells per sample were imaged at maximum width in view of both the nuclei and the PECs bare zone. Autofluorescence background in the anti-HA AF488 channel was subtracted from reporter construct lines by thresholding the laser settings against control non-transgenic cells that were also subject to immunoprobing with primary and secondary antibodies.

Single trophozoite images were subject to deconvolution using Huygens Professional (Scientific Volume Imaging) and further analyzed in FIJI/ImageJ[78,79]. Statistical quantification was performed using the coloc2 plugin to determine signal overlap between channels corresponding to Dextran-TxR and Anti-HA AF488. One hundred Costes’ iterations and a point-spread function set to three were used to calculate Pearson’s, Manders’ 1 and 2 coefficients, and Costes’ p-values on the whole image or specific regions of interest [80]. Two regions of interest were chosen, the bare zone and the peripheral PECs. Co-localization analyses were repeated on ≥ ten cells to determine mean and median statistics. Channels corresponding to anti-HA-AF488, Dextran-Texas Red, and DAPI were pseudo-colored in FIJI/ImageJ. Raw signal overlap statistical data for individual cells belonging to each line, as well as expression quantification summaries are provided as Supplementary Table 6.

### Limited cross-linked co-immunoprecipitation using anti-HA agarose beads

Non-transgenic and transgenic trophozoites expressing reporter constructs were used as baits for crosslinking and protein pulldown via anti-HA agarose beads [16,17,43,51]. Each parasite line was cultured in 1 x T-25 flasks to confluency, followed by harvesting and washing with 20 mL PBS yielding a final volume of *ca.* 2×10^9^ cells. A 500 μL aliquot was set aside for western blot analyses to use as control cell lysate samples. The remainder were fixed in 2.25% formaldehyde in PBS and incubated for 30 minutes at room temperature while shaking. Cells were quenched in 10 mL 0.1M Glycine in PBS for 15 minutes with gentle shaking and resuspended in 5 mL RIPA-SDS lysis buffer solution with 50 mM Tris (pH 7.4), 150 mM NaCl, 1% IGEPAL, 0.5% sodium deoxycholate, 0.1% SDS, and 10 mM EDTA. 100 μL 0.1M phenylmethylsulphonyl fluoride (PMSF; Sigma) and 50 μL protease inhibitor cocktail (Sigma) was also added, and the resulting suspension was sonicated twice using the following settings: 60 pulses, two output control, 30% duty cycle, and 60 pulses, four output control, 40% duty cycle, respectively, with 10-15 seconds break between each cycle. Sonicated samples were incubated at 4°C on a rotating shaker for 2 hours and centrifuged at 14,000 *x g* for 10 minutes at 4°C. Supernatants from each sample were syringe filtered through 0.2 μm Acrodisc MS filters (Pall MS-3301) and resuspended in 5 mL RIPA-Triton X-100 solution containing 50mM Tris (pH 7.4), 150mM NaCl, 1% IGEPAL, 0.5% sodium deoxycholate, 1% Triton X-100, and 10 mM EDTA (Sigma). Forty microlitres of the anti-HA agarose beads slurry (Thermo Fisher) was added to the soluble protein fraction and incubated overnight on a rotating shaker at 4°C. Subsequently, beads were washed thrice with 10 mL Tris-Buffered Saline (TBS) with 0.1% Triton X-100. One hundred microlitre aliquot of the beads was saved for immunoblotting, while the remainder of the sample was subject to mass spectrometry analysis as detailed in section 2.7. For each transgenic line, co-immunoprecipitation and mass-spectrometry experiments were performed in independent biological replicates of two.

### Immuno-Blotting and Coomassie Staining

Proteins from whole cell lysate and bead samples from both transgenic and non-transgenic control were analyzed by SDS-PAGE on a 4%-10% polyacrylamide gel under reducing conditions. Cross-linking was reversed by resuspending bead and whole cell lysate samples in Laemmli loading buffer and 100 mM Dithiothreitol (DTT) (Sigma), then boiling for 5 minutes. Samples were loaded with PageRuler™ Prestained Protein Ladder (10 to 180 kDa; Cat. No. 26616, Thermo Fisher) and run at 160V for 1 hour. Gels were transferred onto nitrocellulose membranes overnight at 4°C and blocked with 5% dry milk/0.05% TWEEN-20 the following day. Immunoprobing was performed using a rat-derived monoclonal anti-HA antibody (dilution 1:1000; Roche) followed by a secondary anti-rat antibody coupled to horseradish peroxidase (HRP) (dilution 1:5000; Southern Biotech). HRP detection was performed by adding Western chemiluminescent substrate (Thermo Fisher) onto the membrane and visualized using an Amersham Imager 600 under the chemiluminescent settings. Polyacrylamide gels were stained with InstantBlue Coomasie Protein Stain (Sigma) for 30 minutes and de-stained with ddH_2_0 while shaking.

### Liquid Chromatography Mass Spectrometry (LC/MS) for quantitative proteomics

Affinity pulldown beads were suspended in 8M Urea / 50 mM Tris-HCl (pH 8). Proteins were reduced at 37°C with DTT 0.1M / 100 mM Tris-HCl (pH 8) and alkylated at 37°C in the dark with IAA 0.5M / 100 mM Tris-HCl (pH 8) for 30 minutes each. Thereafter, the slurry was diluted with four volumes of 20 mM Tris-HCl (pH 8) / 2 mM CaCl_2_ prior to overnight digestion at room temperature with 100 ng sequencing grade trypsin (Promega). Samples were centrifuged in order to extract the peptides in the supernatant. The digests were analyzed by liquid chromatography (LC)-MS/MS using PROXEON coupled to a Q-Exactive mass spectrometer (Thermo Fisher Scientific) with three injections of five microlitre digests. Peptides were trapped on a µPrecolumn C18 PepMap100 (5 μm, 100 Å, 300 μm× 5 mm, Thermo Fisher Scientific, Reinach, Switzerland) and separated by backflush on a C18 column (5 μm, 100 Å, 75 μm×15 cm, C18) by applying a 40 minute gradient of 5% acetonitrile to 40% in water, 0.1% formic acid, at a flow rate of 350 nl/min. The Full Scan method was set with resolution at 70,000, an automatic gain control (AGC) target of 1×10^6^, and a maximum ion injection time of 50 ms. The data-dependent method for precursor ion fragmentation was applied with the following settings: resolution 17,500, AGC of 1×10^5^ maximum ion time of 110 milliseconds, mass isolation window 2 *m/z*, collision energy 27, under fill ratio 1%, charge exclusion of unassigned and 1+ ions, and peptide match preferred, respectively.

MS data were interpreted with MaxQuant (v. 1.6.14.0) against GiardiaDB (v. 47) using the default MaxQuant settings, with an allowed mass deviation for precursor ions of 15 ppm in the first search round, the mass deviation for fragments of 20 ppm, a maximum peptide mass of 5500Da, the match between runs activated with a matching time window of 0.7 min, and the use of non-consecutive fractions for the different pulldowns to prevent over-fitting. Other database search settings were a strict trypsin cleavage rule allowing for 3 missed cleavages, fixed carbamidomethylation of cysteines, variable oxidation of methionines, and acetylation of protein N-termini. Resulting MS hits, including intensity-based absolute quantification (iBAQ) and label-free quantitation (LFQ) values, are provided in Supplementary Table 7.

### In silico evaluation of LC/MS hits

Prior to analyses, contaminants and highly abundant proteins enriched in the non-transgenic control were filtered out from the MS data corresponding to the transgenic lines. Each replicate from a given line was treated independently for initial analyses. MS hits were sorted according to intensity-based absolute quantification (iBAQ) values. iBAQ values measured the sum of peak intensities divided by theoretically observed peptides to quantify the relative molar amount of a given protein in the sample [81]. These were normalized by calculating the relative percent abundance for each protein hit as per the following equation:

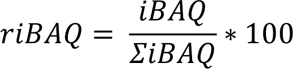

Protein hits from replicate datasets belonging to a given line were intersected with one another, wherein only hits enriched in both replicates were kept for investigation. A high and a low stringency threshold of 1% and 0.01% relative (r)iBAQ, respectively, was applied. All hits above the 0.01% threshold were considered significant molecular interactions and deemed biologically relevant in the context of the bait. Within these, all identified hypothetical proteins were subject to further analyses using HHPRED analyses (https://toolkit.tuebingen.mpg.de/tools/hhpred) and screened for in all previously published *Giardia* co-IP datasets for their functional annotation. Sorted absolute and relative iBAQ hits meeting stringency thresholds are provided in the Supplementary Table 8.

## Supplementary Figures and Tables

**Supplementary Figure 1. Preliminary phylogenetic investigation assessing the classification of ARF homologues in the BaSks.** Classified ARF1 homologues in BaSks and *Anaeramoba* against an ARF backbone [30]. In this, values for supported nodes (SH-aLRT/IQ-TREE-2) have been replaced by symbols: black circles: >95/95, grey circles: >90/90. Node support values are overlaid onto IQ-TREE-2 tree topology.

**Supplementary Figure 2. Preliminary phylogenetic investigation assessing the classification of ARF homologues in the BaSks.** Classified ARF6 homologues in BaSks and *Anaeramoba* against an ARF backbone [30]. In this, values for supported nodes (SH-aLRT/IQ-TREE-2) have been replaced by symbols: black circles: >95/95, grey circles: >90/90. Node support values are overlaid onto IQ-TREE-2 tree topology.

**Supplementary Figure 3. Preliminary phylogenetic investigation assessing the classification of ARF homologues in the BaSks.** Classified ARF1 homologues in BaSks against an ARF backbone [30]. In this, values for supported nodes (SH-aLRT/IQ-TREE-2) have been replaced by symbols: black circles: >95/95, grey circles: >90/90. Node support values are overlaid onto IQ-TREE-2 tree topology.

**Supplementary Figure 4.** Population-level expression analysis of epitope-tagged ARF regulatory system proteins. (**A**) Non-transgenic lines (WB) were used as controls and for the subtraction of background anti-HA AF488 signal. (**B**) GiARF1-HA was expressed in 88% of the screened cells, of which 16% had an overexpression phenotype. (**C**) GiARF1FA-HA was expressed in 89% of the screened cells, of which 13% had an overexpression phenotype. (**D**) GiARF1FB1-HA was expressed in 79% of the screened cells, of which 14% had an overexpression phenotype. (**E**) GiARFGAP1-HA was expressed in 79% of the screened cells, of which 18% had an overexpression phenotype. (**F**) GiCYTHa-HA was expressed in 82% of the screened cells, of which 24% had an overexpression phenotype. (**G**) provides a detailed breakdown of the number of cells that were used for quantification. All scale bars: 5 µm.

**Supplementary Figure 5.** Network analysis of the PECs-associated interacting partners. LC/MS hits enriched in all five Giardia ARF regulatory system proteomic datasets with known PEC localizations and functions were intersected to identify common trends in the molecular machinery involved in vesicle formation and fusion pathways, as well as those with other endo-lysosomal activities. Highly abundant interactions with (r)iBAQ of ≥1% are indicated with solid lines.

**Supplementary Figure 6.** Vector maps for constructs synthesized and presented in this report. Vector maps of plasmids for GiARF1-HA (GL50803_7789), GiARF1FA-HA (GL50803_13930), GiARF1FB1-HA (GL50803_7562), GiARFGAP1-HA (GL50803_22454), and GiCYTHa-HA (GL50803_17912), all cloned into a pPacV-Integ backbone episome.

**Supplementary Table 1.** Table summarizing genome database references.

**Supplementary Table 2.** Table summarizing forward and reverse ARF1, ARF6, ARFGAP, and ARFGEF AMOEBAE searches into the BaSks and *Anaeramoeba*.

**Supplementary Table 3.** Table summarizing reverse BLAST searches of BaSk ARF1 hits.

**Supplementary Table 4.** Table summarizing annotation guide for naming conventions within phylogenetic trees and the corresponding accession available.

**Supplementary Table 5.** Table summarizing forward and reverse oligonucleotide sequences used to clone and express GiARF1, GiARF1FA, GiARF1FB1, GiARFGAP1, and GiCYTHa with C-terminal HA epitope tags.

**Supplementary Table 6.** Quantification of cells expressing recombinant proteins and signal overlap analyses between Dextran-TxR and anti-HA-AF488 for all analyzed Giardia ARF regulatory system proteins.

**Supplementary Table 7.** All raw peptide hits acquired using LC/MS and summary of spectral counts, iBAQ values, and LFQ values for each hit.

**Supplementary Table 8.** Summary of LC/MS hits post decontamination and removal of non-specific enrichments, including overlap analysis for the three ARF paralogue interactomes. Sorting was performed according to highest to lowest protein abundance, as per the relative iBAQ values.

**Supplementary Table 9.** Summary of experimentally validated PEC hits that were enriched with the Giardia ARF regulatory system proteins used as affinity handles in this study. PEC annotation was assigned according to previously published microscopy, co-IP, or yeast-two hybrid evidence.

**Supplementary File** 1. Masked ARF1 alignment used for phylogenetic tree building.

**Supplementary File** 2. Masked ARF6 alignment used for phylogenetic tree building.

**Supplementary File** 3. Sequence of plasmid P91-pE-ARF3-HA (ARF)

**Supplementary File** 4. Sequence of plasmid P92-pE-ARF2-HA

**Supplementary File** 5. Sequence of plasmid P96-pE-ARF1-HA

**Supplementary File** 6. Sequence of plasmid P124-pE-ARF GAP1-HA

**Supplementary File** 7. Sequence of plasmid P125-pE-CYTHa-HA

## Acknowledgements

We thank the Microscopy Imaging Center (MIC) of the University of Bern, particularly Drs. Y. Belyaev and G. Witz for training and advice. We thank the Proteomics and Mass Spectrometry Core Facility (PMSCF) of the Department for Biomedical Research at the University of Bern for generating all proteomics data. All figures assembled using BioRender.

SVP received support and a stipend from a CIHR Doctoral Scholarship (CIHR CGM 175863) and an Alberta Innovates Graduate Student Scholarship. Research in the Dacks lab is funded by NSERC Discovery Grants (RES0043758, RES0046091), awarded to JBD. Research in the Faso lab is funded by Swiss National Science Foundation grant numbers PR00P3_179813, PR00P3_179813/2 and PR00P3_179813/3, awarded to CF.

## Author contributions

SVP: co-designed experimental strategy, generated and analyzed data, co-wrote the manuscript. EK: co-designed experimental strategy, generated and analysed data, co-wrote the manuscript. CW: generated data, supported experimental strategy design. EAB: generated data, supported experimental strategy design. RAK: co-wrote the manuscript and provided deep critical feedback. JBD: provided funding, co-wrote the manuscript. CF: co-designed experimental strategy, generated and analyzed data, provided funding and access to infrastructure, co-wrote the manuscript.

