## supplementary figures 1-6 for "The ARF regulatory GTPase in *Giardia intestinalis* is associated with vesicle formation and membrane fusion machinery at non-canonical endosomal compartments"

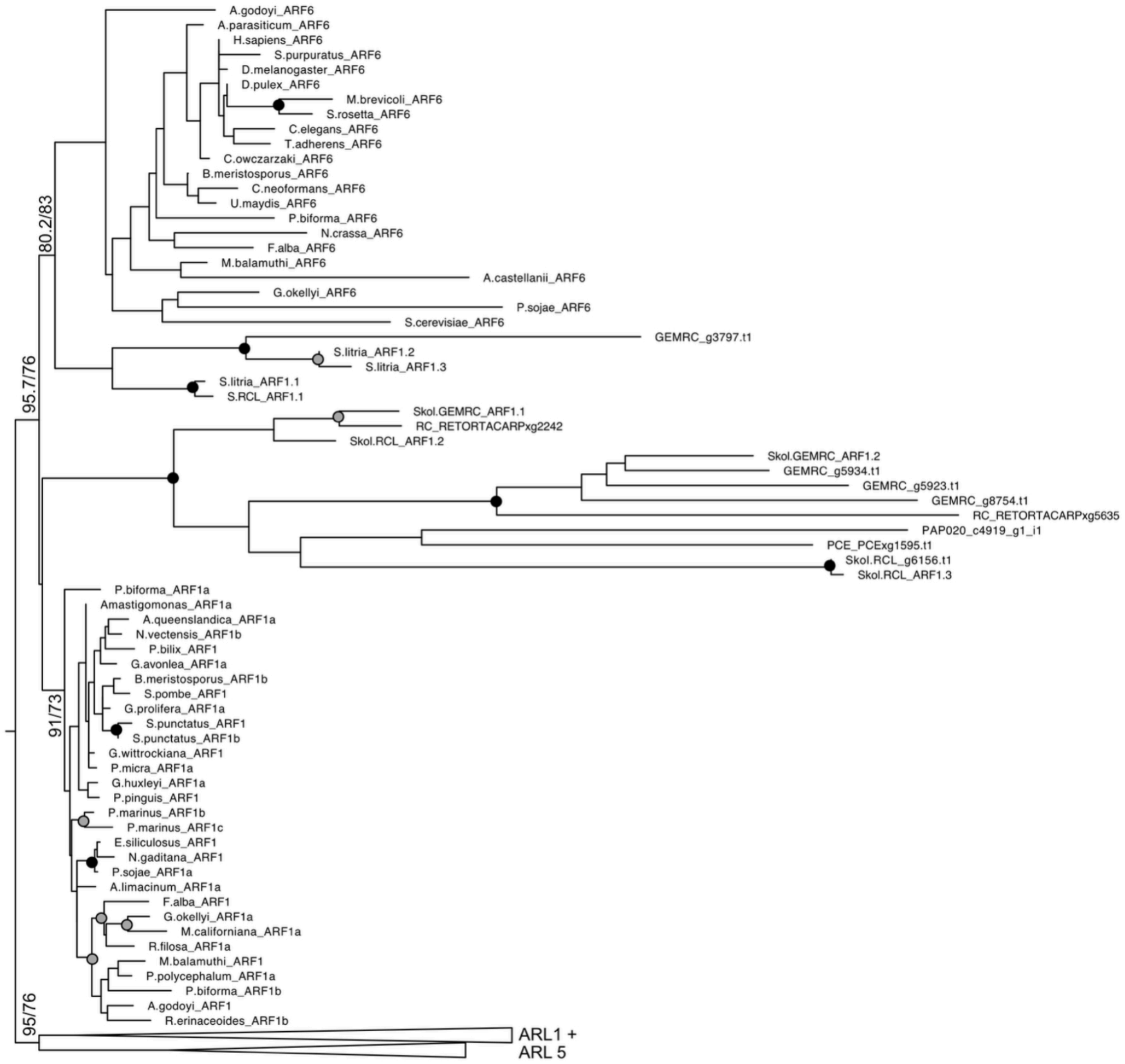

A.godoyi_ARF6
A.parasiticum_ARF6
H.sapiens_ARF6
S.purpuratus_ARF6
D.melanogaster_ARF6
D.pulex_ARF6
M.brevicoli_ARF6
S.rosetta_ARF6
C.elegans_ARF6
T.adherens_ARF6
C.owczarzaki_ARF6
B.meristosporus_ARF6
C.neoformans_ARF6
U.maydis_ARF6
P.biforma_ARF6
N.crassa_ARF6
F.alba_ARF6
M.balamuthi_ARF6
A.castellanii_ARF6
G.okellyi_ARF6
P.sojae_ARF6
S.cerevisiae_ARF6
GEMRC_g3797.t1
S.litria_ARF1.2
S.litria_ARF1.3
S.litria_ARF1.1
S.RCL_ARF1.1
Skol.GEMRC_ARF1.1
RC_RETORTACARPxg2242
Skol.RCL_ARF1.2
Skol.GEMRC_ARF1.2
GEMRC_g5934.t1
GEMRC_g5923.t1
GEMRC_g8754.t1
RC_RETORTACARPxg5635
PAP020_c4919_g1_i1
PCE_PCExg1595.t1
Skol.RCL_g6156.t1
Skol.RCL_ARF1.3
P.biforma_ARF1a
Amastigomonas_ARF1a
A.queenslandica_ARF1a
N.vectensis_ARF1b
P.bilix_ARF1
G.avonlea_ARF1a
B.meristosporus_ARF1b
S.pombe_ARF1
G.prolifera_ARF1a
S.punctatus_ARF1
S.punctatus_ARF1b
G.wittrockiana_ARF1
P.micra_ARF1a
G.huxleyi_ARF1a
P.pinguis_ARF1
P.marinus_ARF1b
P.marinus_ARF1c
E.siliculosus_ARF1
N.gaditana_ARF1
P.sojae_ARF1a
A.limacinum_ARF1a
F.alba_ARF1
G.okellyi_ARF1a
M.californiana_ARF1a
R.filosa_ARF1a
M.balamuthi_ARF1
P.polycephalum_ARF1a
P.biforma_ARF1b
A.godoyi_ARF1
R.erinaceoides_ARF1b
80.2/83
95.7/76
91/73
95/76
ARL1 +
ARL 5

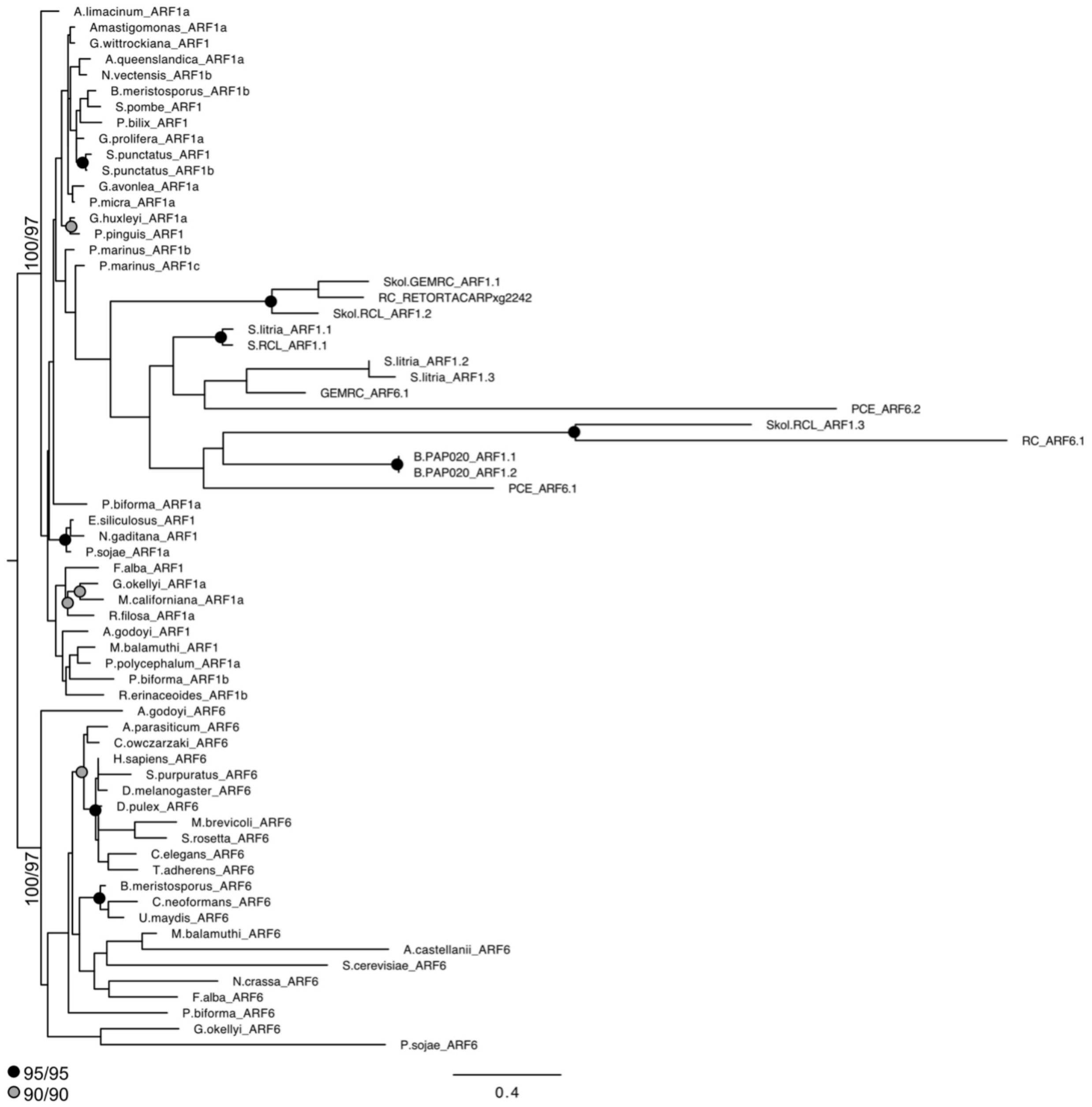
A.limacinum_ARF1a
Amastigomonas_ARF1a
G.wittrockiana_ARF1
A.queenslandica_ARF1a
N.vectensis_ARF1b
B.meristosporus_ARF1b
S.pombe_ARF1
P.bilix_ARF1
G.prolifera_ARF1a
S.punctatus_ARF1
S.punctatus_ARF1b
G.avonlea_ARF1a
P.micra_ARF1a
G.huxleyi_ARF1a
P.pinguis_ARF1
P.marinus_ARF1b
P.marinus_ARF1c
Skol.GEMRC_ARF1.1
RC_RETORTACARPxg2242
Skol.RCL_ARF1.2
S.litria_ARF1.1
S.RCL_ARF1.1
S.litria_ARF1.2
S.litria_ARF1.3
GEMRC_ARF6.1
PCE_ARF6.2
Skol.RCL_ARF1.3
RC_ARF6.1
B.PAP020_ARF1.1
B.PAP020_ARF1.2
PCE_ARF6.1
P.biforma_ARF1a
E.siliculosus_ARF1
N.gaditana_ARF1
P.sojae_ARF1a
F.alba_ARF1
G.okellyi_ARF1a
M.californiana_ARF1a
R.filosa_ARF1a
A.godoyi_ARF1
M.balamuthi_ARF1
P.polycephalum_ARF1a
P.biforma_ARF1b
R.erinaceoides_ARF1b
100/97
A.godoyi_ARF6
A.parasiticum_ARF6
C.owczarzaki_ARF6
H.sapiens_ARF6
S.purpuratus_ARF6
D.melanogaster_ARF6
D.pulex_ARF6
M.brevicoli_ARF6
S.rosetta_ARF6
C.elegans_ARF6
T.adherens_ARF6
B.meristosporus_ARF6
C.neoformans_ARF6
U.maydis_ARF6
M.balamuthi_ARF6
A.castellanii_ARF6
S.cerevisiae_ARF6
N.crassa_ARF6
F.alba_ARF6
P.biforma_ARF6
G.okellyi_ARF6
P.sojae_ARF6
100/97
95/95
90/90
0.4

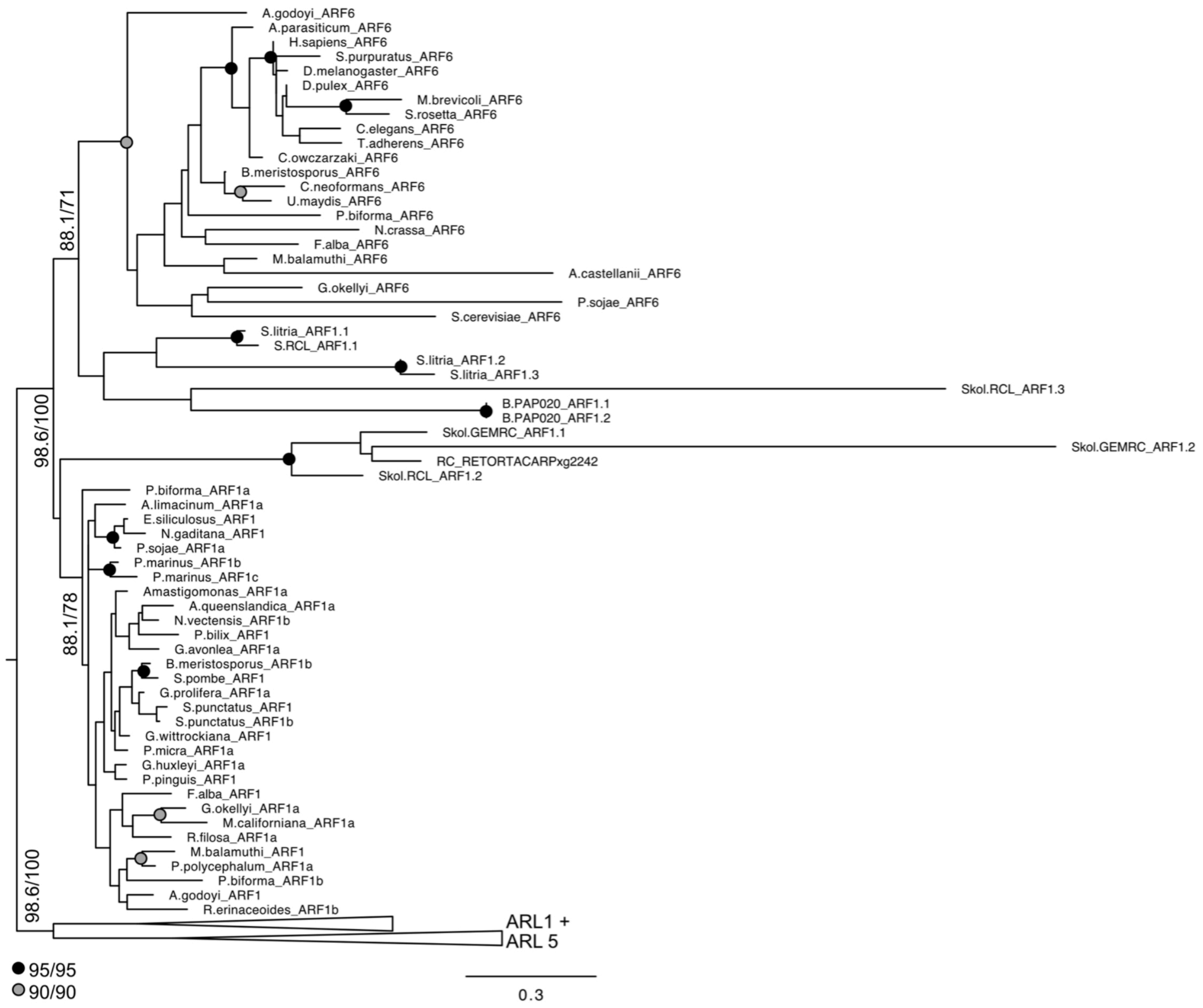
A.godoyi_ARF6
A.parasiticum_ARF6
H.sapiens_ARF6
S.purpuratus_ARF6
D.melanogaster_ARF6
D.pulex_ARF6
M.brevicoli_ARF6
S.rosetta_ARF6
C.elegans_ARF6
T.adherens_ARF6
C.owczarzaki_ARF6
B.meristosporus_ARF6
C.neoformans_ARF6
U.maydis_ARF6
P.biforma_ARF6
N.crassa_ARF6
F.alba_ARF6
M.balamuthi_ARF6
A.castellanii_ARF6
G.okellyi_ARF6
P.sojae_ARF6
S.cerevisiae_ARF6
S.litria_ARF1.1
S.RCL_ARF1.1
S.litria_ARF1.2
S.litria_ARF1.3
Skol.RCL_ARF1.3
B.PAP020_ARF1.1
B.PAP020_ARF1.2
Skol.GEMRC_ARF1.1
Skol.GEMRC_ARF1.2
RC_RETORTACARPxg2242
Skol.RCL_ARF1.2
P.biforma_ARF1a
A.limacinum_ARF1a
E.siliculosus_ARF1
N.gaditana_ARF1
P.sojae_ARF1a
P.marinus_ARF1b
P.marinus_ARF1c
Amastigomonas_ARF1a
A.queenslandica_ARF1a
N.vectensis_ARF1b
P.bilix_ARF1
G.avonlea_ARF1a
B.meristosporus_ARF1b
S.pombe_ARF1
G.prolifera_ARF1a
S.punctatus_ARF1
S.punctatus_ARF1b
G.wittrockiana_ARF1
P.micra_ARF1a
G.huxleyi_ARF1a
P.pinguis_ARF1
F.alba_ARF1
G.okellyi_ARF1a
M.californiana_ARF1a
R.filosa_ARF1a
M.balamuthi_ARF1
P.polycephalum_ARF1a
P.biforma_ARF1b
A.godoyi_ARF1
R.erinaceoides_ARF1b
ARL1 +
ARL 5
88.1/71
98.6/100
88.1/78
98.6/100
95/95
90/90
0.3

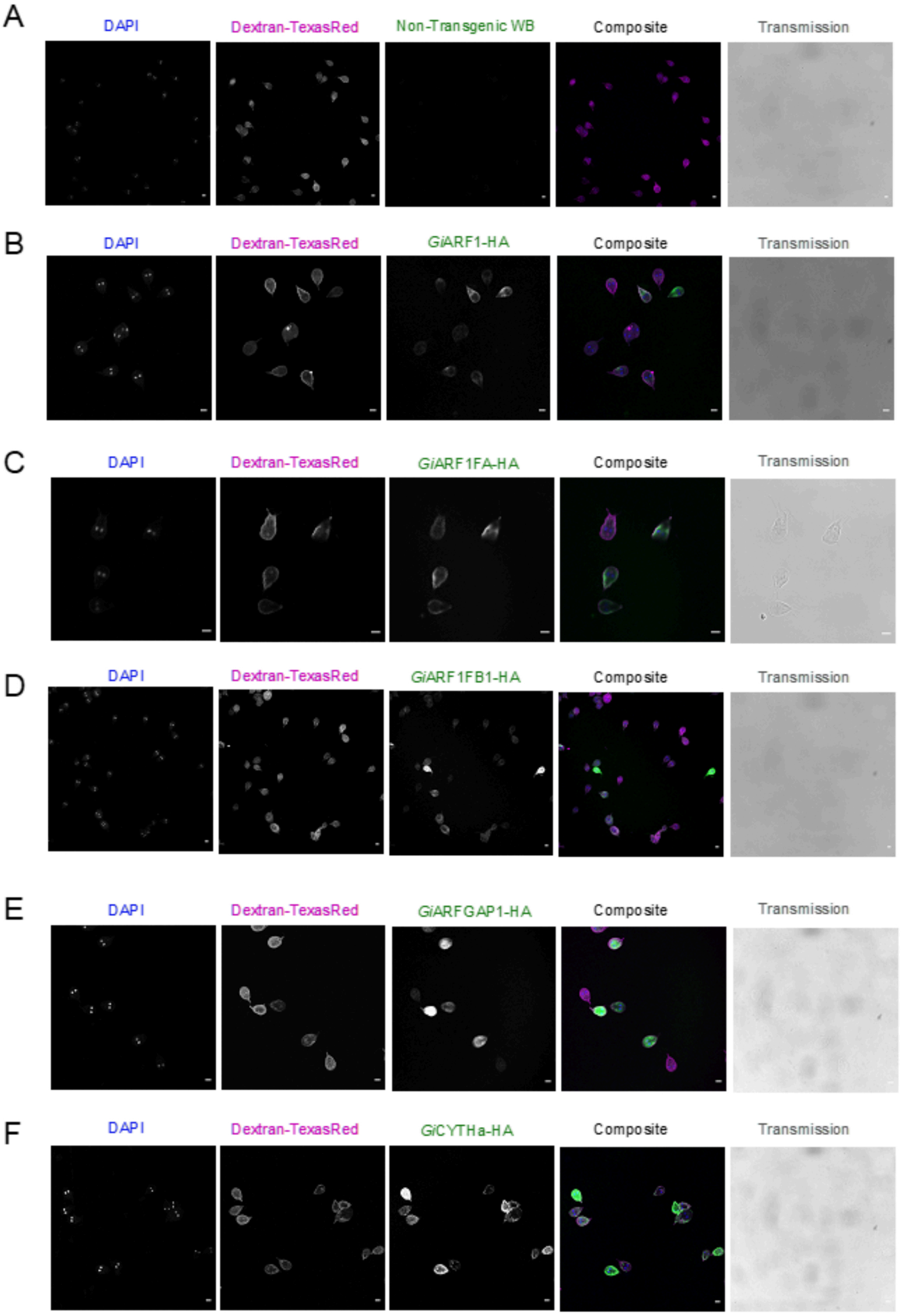


G

| Giardia cell line | Transgenic construct | Constitutive or Inducible expression | Total cell number counted | Total cells expressing | Total cells overexpressing | Percentage cells expressing | Percentage cells overexpressing |
|---|---|---|---|---|---|---|---|
| G96 | pE_*Gi*ARF1_HA (GL50803_7789) | constitutive | 154 | 138 | 25 | 88 | 16 |
| G91 | pE_*Gi*ARF1FA_HA (GL50803_13930) | constitutive | 189 | 162 | 24 | 89 | 13 |
| G92 | pE_*Gi*ARF1FB1_HA (GL50803_7562) | constitutive | 471 | 370 | 68 | 79 | 14 |
| G124 | pE_*Gi*ARFGAP1_HA (GL50803_2834) | constitutive | 461 | 365 | 84 | 79 | 18 |
| G125 | pE_*Gi*CYTHa_HA (GL50803_17192) | constitutive | 380 | 313 | 93 | 82 | 24 |

**Supplementary Figure 4**. **Population-level expression analysis of epitope-tagged ARF regulatory system proteins**. (**A**) Non-transgenic lines (WB) were used as controls and for the subtraction of background anti-HA AF488 signal. (**B**) *Gi*ARF1-HA was expressed in 88% of the screened cells, of which 16% had an overexpression phenotype. (**C**) *Gi*ARF1FA-HA was expressed in 89% of the screened cells, of which 13% had an overexpression phenotype. (**D**) *Gi*ARF1FB1-HA was expressed in 79% of the screened cells, of which 14% had an overexpression phenotype. (**E**) *Gi*ARFGAP1-HA was expressed in 79% of the screened cells, of which 18% had an overexpression phenotype. (**F**) *Gi*CYTHa-HA was expressed in 82% of the screened cells, of which 24% had an overexpression phenotype. (**G**) provides a detailed breakdown of the number of cells that were used for quantification. All scale bars: 5 µm.

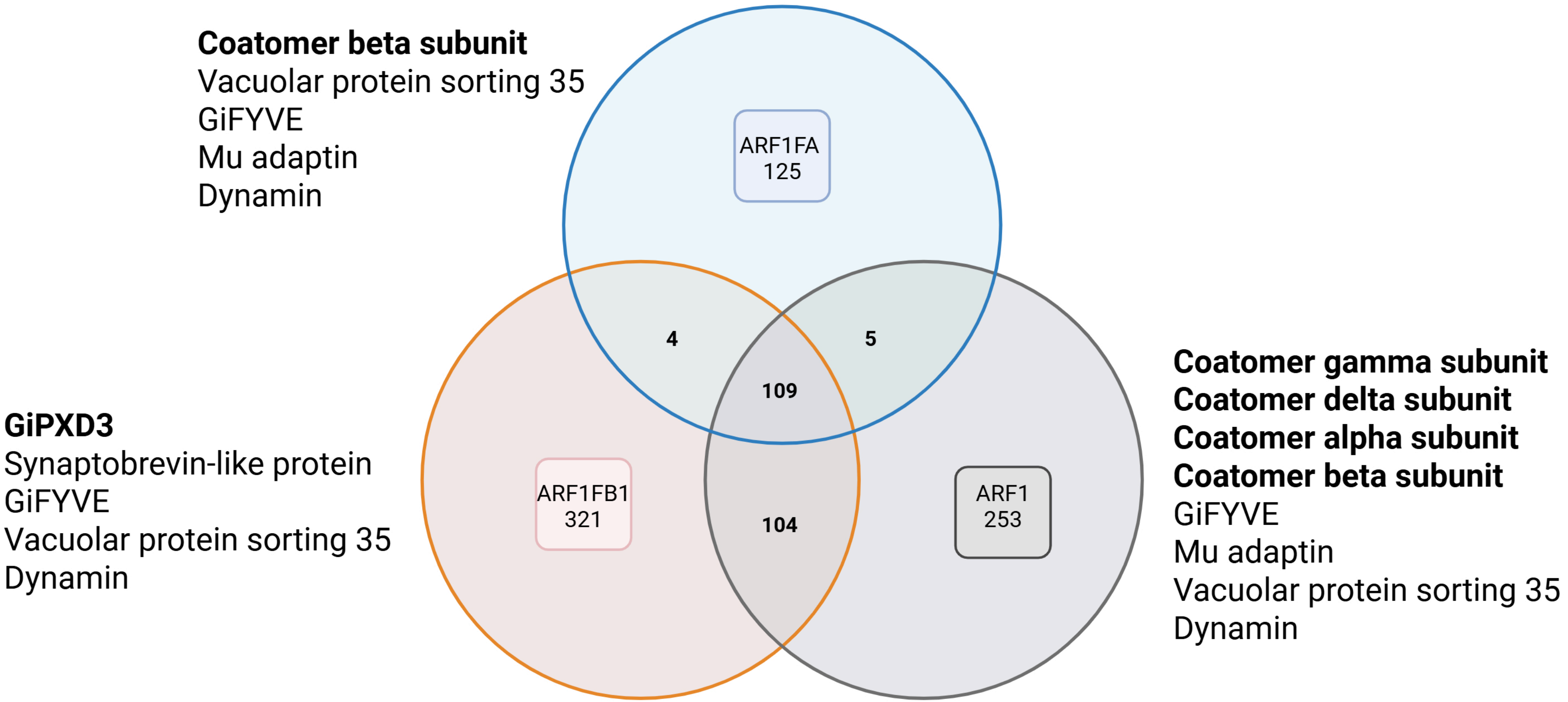


**Supplementary figure 5.** Venn diagrams for protein interactome datasets of the three ARF paralogues. A core of 109 proteins is shared amongst the three curated datasets. For a complete list of proteins found in the intersects, including their ORF number and putative function based on available annotations, see supplementary table 4 . Known and reported PEC-associated proteins are listed for each dataset are summarised in supplementary table 5.

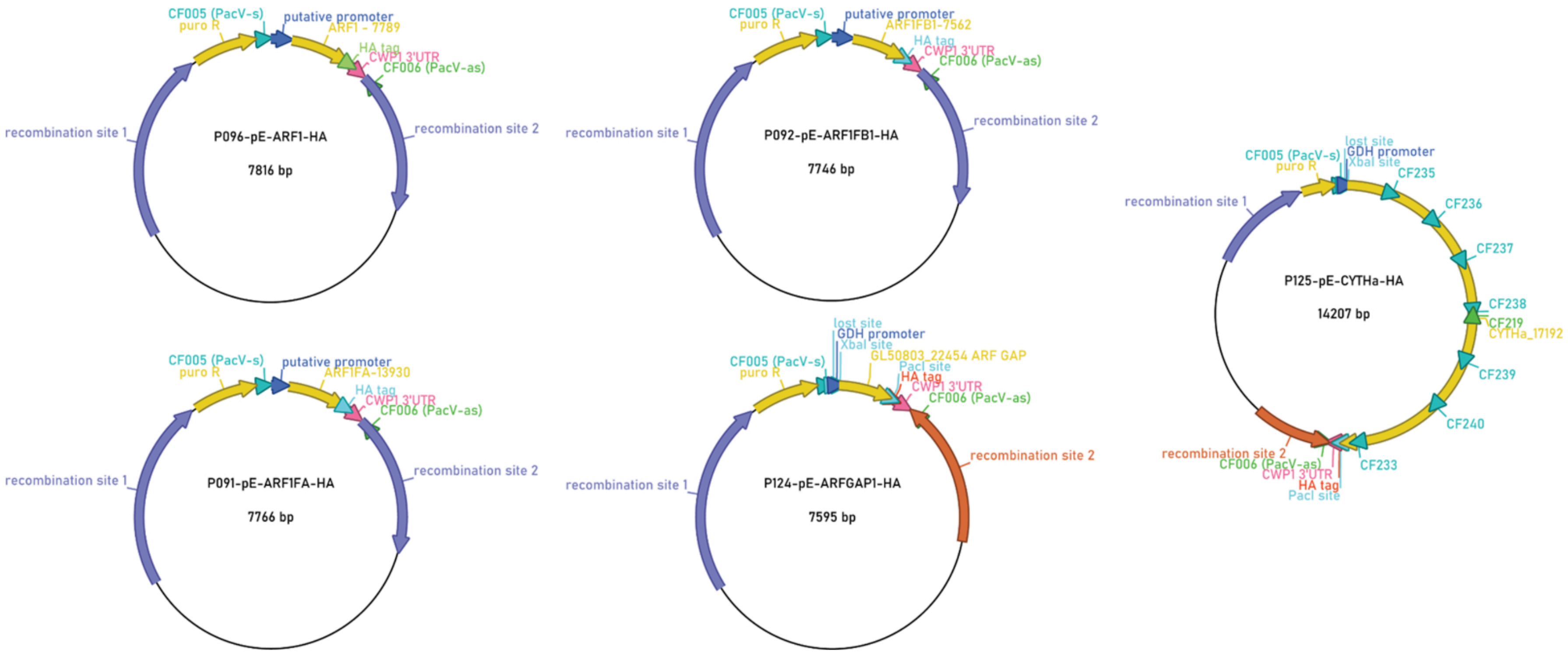


**Supplementary Figure 6**. Vector maps for constructs synthesized and presented in this report. Vector maps of plasmids for GiARF1-HA (GL50803_7789), GiARF1FA-HA (GL50803_13930), GiARF1FB1-HA (GL50803_7562), GiARFGAP1-HA (GL50803_22454), and GiCYTHa-HA (GL50803_17912), all cloned into a pPacV-Integ backbone episome. Sequence information is provided as a .txt file for each construct sh
